# Therapeutic AIBP expression contributes to synaptic preservation and vision restoration against glaucomatous neuroinflammation

**DOI:** 10.64898/2026.10.10.758145

**Authors:** Tonking Bastola, Sinwoo Hwang, Ziyao Shen, Soo-Ho Choi, Seunghwan Choi, Esther Jang, Muna Poudel, Keun-Young Kim, Yury I. Miller, Robert N. Weinreb, Won-Kyu Ju

**Author notes:** These authors contributed equally (Tonking Bastola, Sinwoo Hwang, Ziyao Shen). Correspondence to Won-Kyu Ju, Gleiberman Center for Glaucoma Research, Viterbi Family Vision Research Center and Shiley Eye Institute, The Viterbi Family Department of Ophthalmology, University of California San Diego, 9500 Gilman Drive, La Jolla, CA 92093, USA.

## Abstract

Lowering intraocular pressure (IOP) is the only approved treatment for glaucoma, yet retinal ganglion cell (RGC) degeneration and vision loss often progress despite reducing IOP. Apolipoprotein A-I binding protein (AIBP) stabilizes ATP-binding cassette subfamily A member 1 (ABCA1) and promotes cholesterol efflux, thus controlling cholesterol-rich lipid rafts and restraining lipid raft-dependent Toll-like receptor 4 (TLR4) inflammatory signaling. Here we show that AAV2-mediated expression of human AIBP (hAIBP) preserves inner retinal synaptic integrity and visual function in experimental glaucoma without altering IOP. In DBA/2J mice, a single intravitreal injection of AAV2-hAIBP at 5 months protected RGCs and preserved pattern electroretinogram and pattern visual evoked potential responses at 10 months, even though IOP was indistinguishable from empty-vector controls. AAV2-hAIBP preserved CaMKII and synapsin phosphorylation, restored synaptophysin and PSD95 in the inner plexiform layer, and reduced the expression of TLR4, complement C1q, IBA1, IL-1β, and IL-6 induction in Müller glia and microglia. Comparable protection was observed in microbead-induced ocular hypertension in C57BL/6 mice, both when the AAV2-hAIBP was delivered before microbead injection and when it was delivered after IOP elevation was established. Elevated IOP reduced endogenous retinal AIBP and ABCA1 expression, and AAV2-hAIBP restored both. Protection required the TLR4-binding domain, as an AIBP mutant lacking this domain failed to preserve RGCs. Conversely, *Apoa1bp^−/−^*retinas exhibited increased CaMKII phosphorylation and reduced synaptophysin in the absence of elevated IOP. These findings identify synaptic preservation through the AIBP–TLR4 axis as an IOP-independent therapeutic mechanism and support AAV2-hAIBP as a candidate gene therapy for glaucoma.

**Graphic Abstract:** 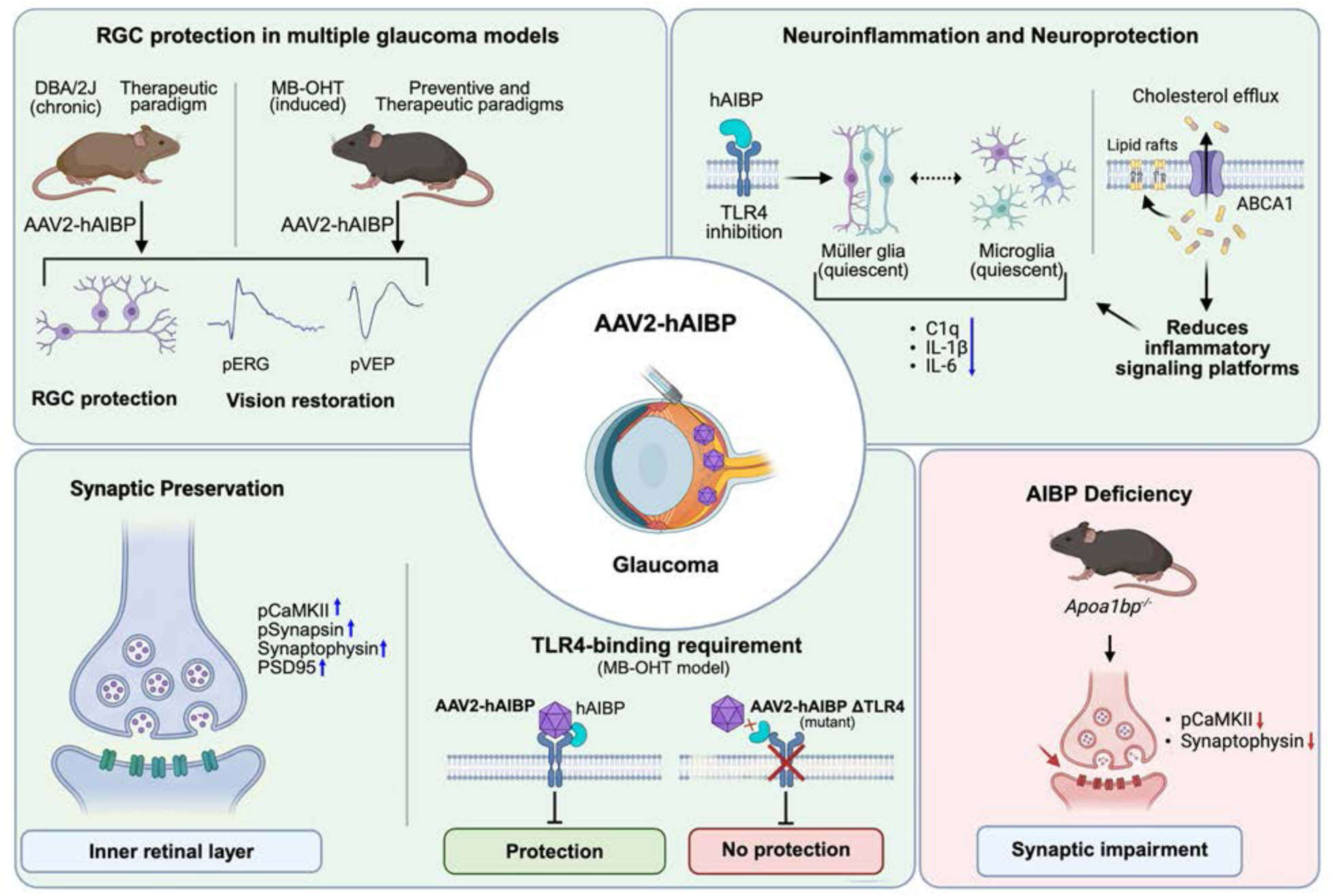

AAV2-hAIBP expression protects RGCs and preserves visual function in chronic DBA/2J (therapeutic mode) and microbead-induced ocular hypertension (preventive and therapeutic modes) without lowering IOP. Retinal hAIBP expression promotes ABCA1-dependent cholesterol efflux, limits lipid raft–dependent TLR4 signaling, keeps Müller glia and microglia quiescent, and reduces C1q, IL-1β, and IL-6. In the IPL, retinal hAIBP expression normalizes CaMKII and synapsin phosphorylation and restores synaptophysin and PSD95. Protection requires the TLR4-binding domain, and *Apoa1bp^−/−^*retinas show synaptic impairment without elevated IOP.

## INTRODUCTION

Glaucoma is a multifactorial neurodegenerative disease of the optic nerve and retinal ganglion cells (RGCs), leading to vision loss.^1,2^ Intraocular pressure (IOP) remains the principal modifiable risk factor, and all currently approved therapies lower IOP. Nonetheless, a substantial proportion of glaucoma patients continue to exhibit progressive RGC loss and visual field deterioration despite IOP being maintained within the target range.^3^ This suggests that IOP-independent mechanisms, such as neuroinflammation,^4,5^ contribute to glaucomatous neurodegeneration. Neuroprotective strategies that preserve RGC somata and axons independently of IOP, therefore, represent an unmet clinical need.

Glia-driven neuroinflammation is a well-known IOP-independent mechanism.^5,6^ Toll-like receptor 4 (TLR4) signaling is upregulated in Müller glia and microglia in glaucomatous human and mouse retina^7^; TLR4 polymorphisms are associated with glaucoma risk, and genetic deletion or pharmacological inhibition of TLR4 reduces RGC death in experimental glaucoma.^8,9^ Direct TLR4 antagonism has nevertheless failed to translate clinically, motivating approaches that target the receptor’s regulatory context rather than the receptor itself. TLR4 activation requires ligand-induced dimerization within cholesterol-rich lipid rafts, so raft cholesterol content is an upstream, tractable control point for the downstream inflammatory program.^10,11^

Apolipoprotein A-I binding protein (AIBP; gene *Apoa1bp*, also known as *NAXE*) precisely regulates this step. AIBP accelerates cholesterol efflux via the transporter ATP-binding cassette subfamily A member 1 (ABCA1), depleting membrane rafts and preventing TLR4 dimerization.^12–14^ Critically, AIBP binds TLR4 directly and acts selectively on inflammatory or cholesterol-loaded cells while leaving resting cells unaffected, distinguishing it from global cholesterol-lowering agents and suggesting a favorable therapeutic index.^13^ AIBP administration reduces neuroinflammation and reverses established pain states in the spinal cord, as well as limiting microglial activation in a model of Alzheimer’s disease.^13,15,16^

In the eye, AIBP and ABCA1 expression decrease while TLR4, interleukin 1β (IL-1β), and retinal cholesterol increase in patients with glaucoma and in mouse models; AIBP knockout (*Apoa1bp^−/−^*) mice exhibit Müller glia activation, mitochondrial dysfunction, and heightened RGC vulnerability to elevated IOP.^7,17^ We recently showed that restoring AIBP with a single intravitreal injection of AAV protects RGCs and their axons and improves visual function in experimental glaucoma, whereas RGC-specific AIBP knockdown worsens RGC function.^7^

Three translational questions remain open to be addressed. First, RGC dendritic and synaptic remodeling precedes soma loss in glaucoma,^18–21^and complement-dependent synapse elimination is an early event in this and other neurodegenerative diseases.^22,23^ Yet, whether AIBP preserves synaptic integrity, rather than merely delaying cell death, remains unknown. Second, prior work delivered the adeno-associated virus (AAV) vector before or at the onset of injury^7^, whereas clinical use would necessarily begin be initiated after diagnosis of the disease. Third, although AIBP has multiple reported activities, including cholesterol efflux-independent effects on raft organization and an intracellular mitochondrial role,^7,24,25^ whether retinal neuroprotection requires its TLR4-binding function specifically has not been tested.

In this study, we used two mechanistically distinct glaucoma models of ocular hypertension (OHT) models of glaucoma to address these questions. We observed that AAV2-mediated hAIBP administration promoted RGC survival, restored pre- and postsynaptic protein content, and preserved retina-to-cortex signal transmission without altering IOP. Furthermore, AIBP administration normalizes calcium/calmodulin-dependent protein kinase II (CaMKII) and synapsin phosphorylation and suppresses TLR4 activation, complement C1q expression, and cytokine induction in glaucomatous retinas. In addition, AIBP deficiency increased CaMKII phosphorylation but reduced retinal synaptophysin expression. Importantly, AIBP administration was effective when delivered after IOP elevation was established (therapeutic mode), and deletion of the TLR4-binding domain abolished AIBP-mediated neuroprotective effects. Together, these findings define synaptic preservation through the AIBP–TLR4 axis as an IOP-independent therapeutic mechanism in glaucoma.

## RESULTS

### Therapeutic mode of AIBP expression protects RGCs and preserves visual function in glaucomatous D2 mice

Because RGCs in 5-month-old DBA/2J (D2) mice are highly susceptible to IOP-related stress before detectable somal or axonal loss,^26,27^ we selected this age as the therapeutic window for intervention. We first characterized visual function in D2 mice and non-glaucomatous DBA/2J-*Gpnmb*^+^ (D2G) controls (Figure 1). At 5 months of age, pattern electroretinogram (pERG) amplitude, pattern visual evoked potential (pVEP) amplitude, and pVEP latency in D2 mice were indistinguishable from those of age-matched control D2G mice (Figure 1A–C), indicating that RGC and post-retinal function remain intact at this age. By 10 months, D2 mice had developed a profound loss of pERG amplitude (approximately 80% reduction, *p* < 0.0001), which reports RGC-driven inner retinal activity; a significant reduction in pVEP amplitude (*p* < 0.05), and a marked prolongation of pVEP latency (*p* < 0.01), which additionally interrogates signal transmission along the optic nerve to the visual cortex, compared to D2G controls (Figure 1D–F). We therefore confirmed vector delivery at 5 months of age and outcome assessment at 10 months. Given that AAV2-GFP transduced 19.5 ± 7.9% of Brn3a-positive RGCs in glaucomatous D2 mice^28^, we next evaluated the transduction efficiency of AAV2-hAIBP-GFP in the retina of 3-month-old C57BL/6J mice at one and seven months post-intravitreal injection (Figure S1 and Table S1). Among RBPMS-positive RGCs, AAV2-hAIBP-GFP transduction efficiency was 19.2 ± 3.1% at one month and increased to 55.2 ± 1.3% at seven months post-injection (Table S2).

**Figure 1.**
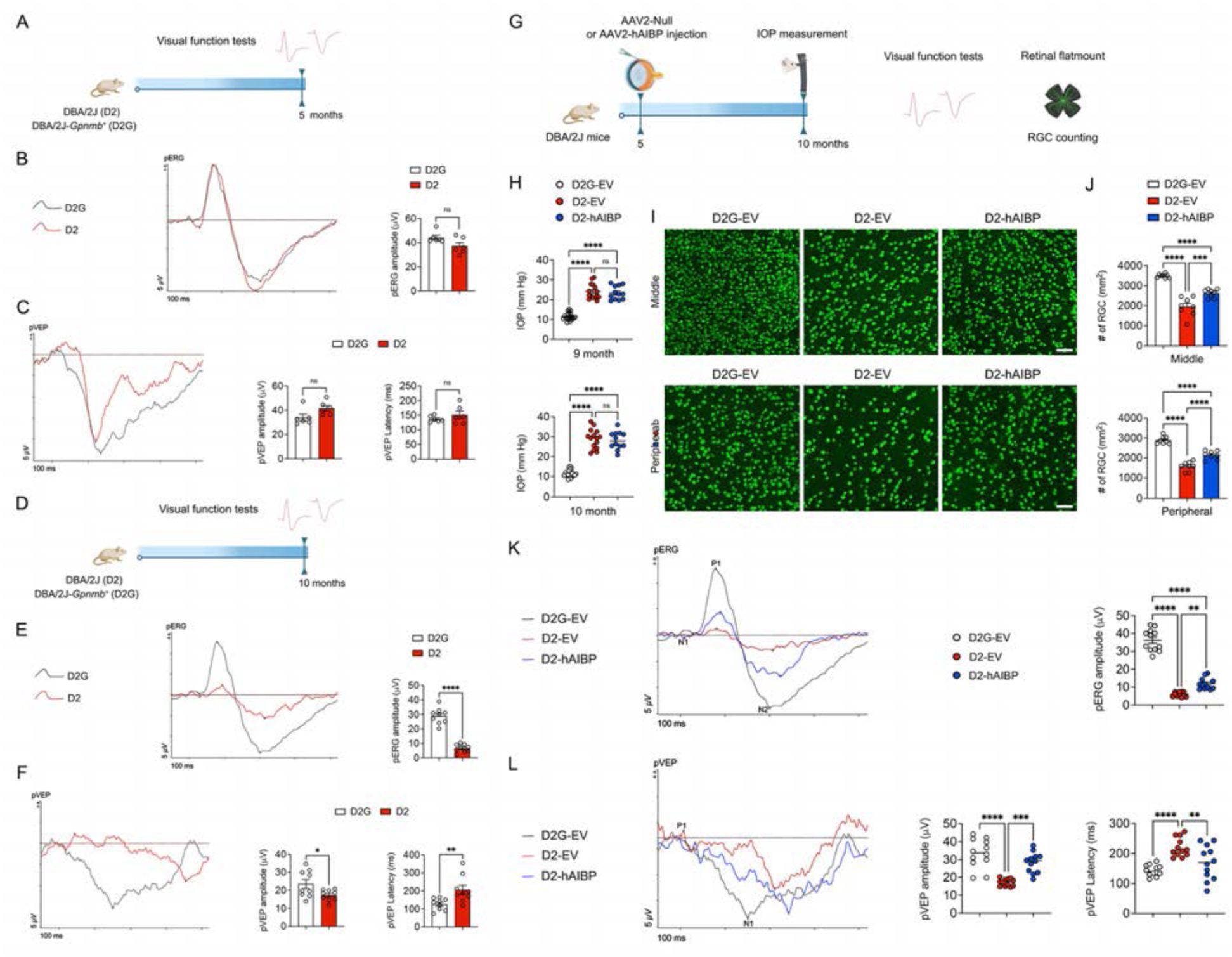
Therapeutic mode of AIBP expression protects RGCs and preserves visual function in glaucomatous D2 mice. (**A**) Experimental schematic and timeline of vision test and data analysis in 5-month-old D2 mice. (**B**) Representative recording graphs for pERG tests and quantitative analysis of pERG amplitude (*n* = 7 mice per group). (**C**) Representative recording graphs for pVEP tests and quantitative analysis of pVEP amplitude and latency (*n* = 7 mice per group). (**D**) Experimental schematic and timeline of vision test and data analysis in 10-month-old D2 mice. (**E**) Representative recording graphs for pERG tests and quantitative analysis of pERG amplitude (*n* = 7 mice per group). (**F**) Representative recording graphs for pVEP tests and quantitative analysis of pVEP amplitude and latency (*n* = 7 mice per group). (**G**) Experimental schematic and timeline of AAV2-hAIBP injection, IOP measurement, vision test, tissue collection, and data analysis in glaucomatous D2 mice. (**H**) IOP measurement in 9 or 10-month-old D2-*Gpnmb^+^*-EV, D2-EV, and D2-hAIBP mice (*n* = 24 mice per group). (**I**) Representative retinal wholemount images for RBPMS (green)-positive RGCs in the middle and peripheral retina. (**J**) Quantitative analysis of RGC numbers in the middle and peripheral areas of the retina (*n* = 12 retina wholemounts from 12 mice for age-matched control Null-D2-*Gpnmb^+^* mice, *n* = 12 retina wholemounts from 12 mice for glaucomatous Null-D2 mice and *n* = 15 retina wholemounts from 15 mice for glaucomatous AIBP-D2 mice). (**K**) Representative recording graphs for pERG tests and quantitative analysis of pERG amplitude (*n* = 7 mice per group). (**L**) Representative recording graphs for pVEP tests and quantitative analysis of pVEP amplitude and latency (*n* = 7 mice per group). Error bars represent SEM. Statistical analysis was performed using one-way ANOVA and Tukey’s multiple comparisons test. \*\**P* < 0.01, \*\*\**P* < 0.001, and \*\*\*\**P* < 0.0001. Scale bar, 50 μm. Panels (A, D, and G) include graphical elements created by BioRender (www.biorender.com).

To test whether restoring human AIBP (hAIBP) in the retina in therapeutic mode is protective in glaucomatous D2 mice, we injected AAV serotype 2 (AAV2) encoding hAIBP (AAV2-hAIBP) or an empty vector (AAV2-EV) intravitreally into 5-month-old D2 mice, before the onset of appreciable IOP elevation in this strain. Age-matched D2G mice injected with AAV2-EV served as normotensive controls. IOP, visual function, and RGC survival were assessed at 10 months of age (Figure 1G). Since the peak IOP was ∼20 mmHg in 9- to 10-month-old D2 mice^29^, IOP was comparable between D2-EV and D2-hAIBP eyes at 9 months (Figure 1H, upper; Table S3). By 10 months, both D2 groups had developed marked OHT relative to D2G-EV controls (D2-EV and D2-hAIBP vs. D2G-EV, *p* < 0.0001), with no difference between D2-EV and D2-hAIBP (Figure 1H, lower; Table S3). AAV2-hAIBP therefore does not act by modifying aqueous outflow or IOP, allowing any downstream benefit to be attributed to a pressure-independent, neuroprotective mechanism.

RGC density was quantified in the RGC marker RNA-binding protein with multiple splicing (RBPMS)-labeled retinal flatmounts in the middle and peripheral retina (Figure 1I; Table S3). Glaucomatous D2-EV retinas lost approximately 45% of RGCs in the middle retina and 45% in the periphery relative to D2G-EV (*p* < 0.0001 for both). hAIBP expression significantly attenuated this loss in both regions (middle, *p* < 0.001; peripheral, *p* < 0.0001 vs. D2-EV), although RGC density in D2-hAIBP retinas remained below that of normotensive D2G-EV controls (*p* < 0.0001; Figure 1J and Table S3). Preserved RGC number was accompanied by preserved RGC function. pERG showed a near-complete collapse of the P1–N2 response in D2-EV eyes (approximately 85% reduction vs. D2G-EV, *p* < 0.0001), which was partially but significantly restored by hAIBP (p < 0.01 vs. D2-EV; Figure 1K). pVEP was likewise reduced in amplitude (*p* < 0.0001) and delayed in latency (*p* < 0.0001) in D2-EV mice. In D2-hAIBP eyes, pVEP amplitude was restored to levels indistinguishable from normotensive controls (*p* < 0.001 vs. D2-EV), and latency was significantly shortened (*p* < 0.01 vs. D2-EV; Figure 1K). Thus, therapeutic administration of hAIBP preserves both the RGC soma population and the functional integrity of the retina–brain pathway despite sustained elevation in IOP.

### hAIBP attenuates TLR4-associated neuroinflammation and glial activation in glaucomatous D2 retina

AIBP is an established regulator of TLR4 lipid-raft signaling.^13,30^ We next examined neuroinflammatory activation in glaucomatous retina. TLR4 immunoreactivity, which co-localized extensively with the Müller glia marker glutamine synthetase (GS), was increased in the inner retina, including the inner plexiform layer (IPL) and ganglion cell layer (GCL), of D2-EV mice and was reduced to control levels in D2-hAIBP retinas (inner retina and IPL, *p* < 0.001; GCL, *p* < 0.01 vs. D2-EV; Figure 2A and B). Downstream inflammatory effectors followed the same pattern. Complement C1q (C1q), a mediator of synapse elimination, was strongly upregulated in the IPL and GCL of D2-EV retinas (*p* < 0.0001) and was significantly reduced by hAIBP (IPL, *p* < 0.05; GCL, *p* < 0.01), although it remained above normotensive levels (p < 0.0001; Figure 2C and D). IBA1-positive microglia showed increased signal intensity in the D2-EV IPL (*p* < 0.0001), which was attenuated by hAIBP (*p* < 0.05 vs. D2-EV; Figure 2E and F). The pro-inflammatory cytokines IL-1β and IL-6, both largely co-localizing with GS-positive Müller glia, were elevated across the inner retina, IPL, and GCL of D2-EV mice and were significantly lowered in D2-hAIBP retinas in every compartment examined (IL-1β: *p* < 0.001, *p* < 0.05, and *p* < 0.001; IL-6: *p* < 0.01 and *p* < 0.01; Figure 2G–J). These findings identify suppression of glial TLR4-driven inflammation and complement induction as a plausible mechanism linking AIBP to retinal synapse and RGC preservation.

**Figure 2.**
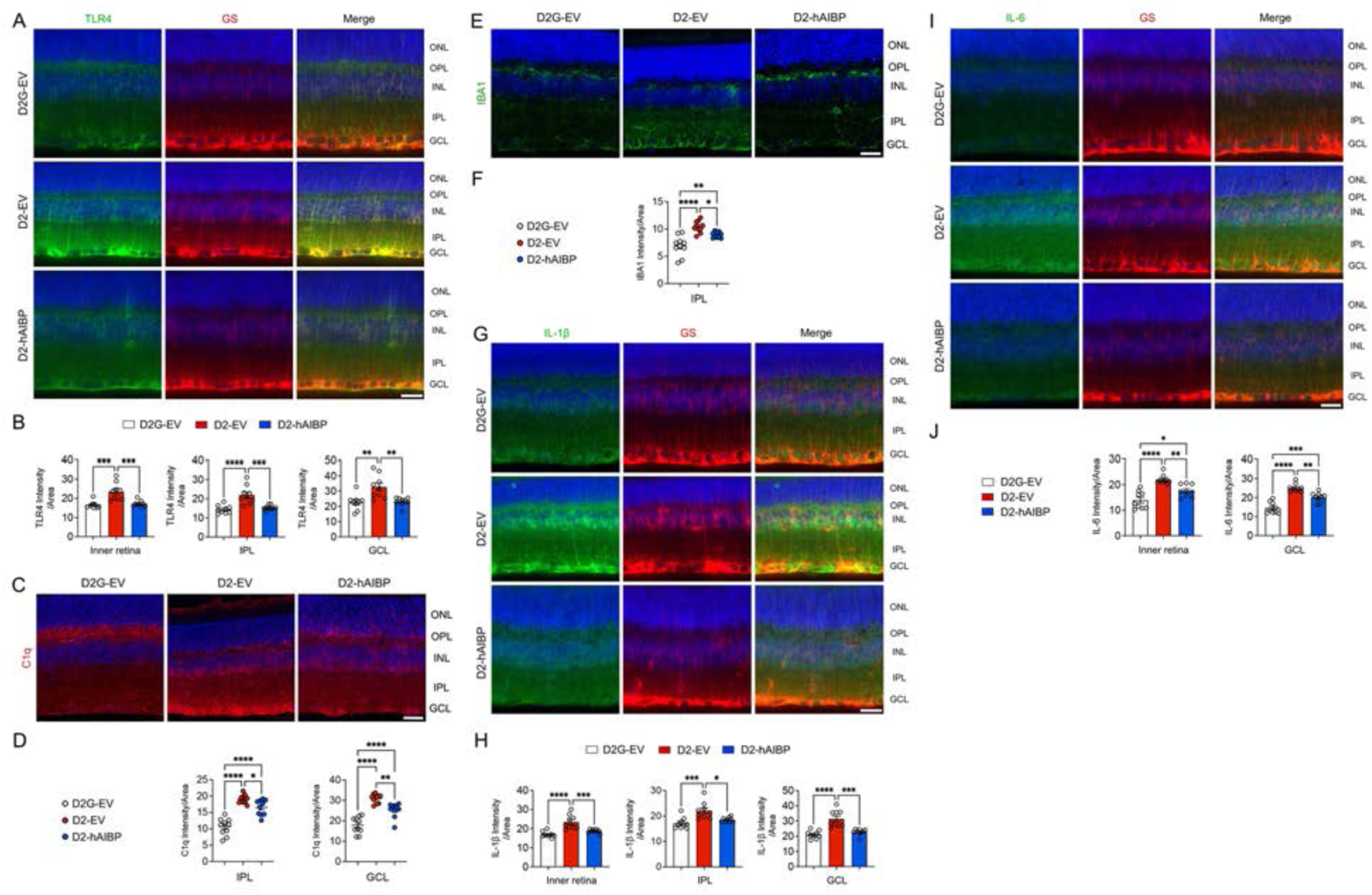
hAIBP attenuates TLR4-associated neuroinflammation and glial activation in glaucomatous D2 retina. (**A**) Representative retinal images for TLR4 (green) and GS (red) immunoreactivity. (**B**) Quantitative fluorescent intensity of TLR4 in the inner layer, IPL, and GCL of the glaucomatous retina (*n* = 5 retina sections from 3 mice per group). (**C**) Representative retinal images for C1q (red) immunoreactivity. (**D**) Quantitative fluorescent intensity of C1q in the IPL and GCL of the glaucomatous retina (*n* = 6 retina sections from 3 mice per group). (**E**) Representative retinal images for IBA1 (green) immunoreactivity. (**F**) Quantitative fluorescent intensity of IBA1 in the IPL of the glaucomatous retina (*n* = 6 retina sections from 3 mice per group). (**G**) Representative retina images for IL-1β (green) and GS (red) immunoreactivity. (**I**) Quantitative fluorescent intensity of IL-1β immunoreactivity in the inner layer, IPL, and GCL of the glaucomatous retina (*n* = 6 retina sections from 3 mice per group). (**G**) Representative retina images for IL-6 (green) and GS (red) immunoreactivity. (**I**) Quantitative fluorescent intensity of IL-6 immunoreactivity in the inner layer and GCL of the glaucomatous retina (*n* = 6 retina sections from 3 mice per group). Error bars represent SEM. Statistical analysis was performed using one-way ANOVA and Tukey’s multiple comparisons test. \**P* < 0.05, \*\**P* < 0.01, \*\*\**P* < 0.001, and \*\*\*\**P* < 0.0001. Scale bars, 20 μm.

### AIBP normalizes CaMKII and synapsin phosphorylation and preserves synaptic proteins in the glaucomatous D2 inner retina

Given that pVEP amplitude was preserved to a greater extent than RGC number, and IBA1 and C1q expression was significantly reduced in the IPL in glaucomatous D2 retina following hAIBP administration, we examined whether hAIBP preserves synaptic integrity in the IPL. We first explored the levels of phosphorylated CaMKII (p-CaMKII), a central regulator of synaptic plasticity, and phosphorylation of synapsin at Ser9 (p-synapsin), a downstream target of CaMKII. Upon Ca²⁺/calmodulin binding, CaMKII undergoes autophosphorylation at Thr286, which renders the kinase autonomously active after Ca²⁺/CaM dissociation. Active CaMKII then phosphorylates downstream synaptic substrates, including synapsins, thereby releasing vesicles from the actin-bound reserve pool and promoting their mobilization and subsequent neurotransmitter release.^31–34^ Immunoblotting of retinal lysates revealed a marked increase in p-CaMKII and p-Synapsin in D2-EV retinas compared to D2G-EV (*p* < 0.01 and *p* < 0.05, respectively), with no change in total synapsin, indicating a shift in phosphorylation state rather than in protein abundance. hAIBP administration normalized both p-CaMKII/actin and p-synapsin/synapsin to control levels (*p* < 0.01 vs. D2-EV; not significant vs. D2G-EV; Figure 3A). Immunostaining localized these changes to the IPL and GCL. p-CaMKII and p-Synapsin immunoreactivity were strongly elevated in the IPL of D2-EV retinas (*p* < 0.0001 vs. D2G-EV) but were significantly reduced by hAIBP administration (p-CaMKII, *p* < 0.001; p-Synapsin, p < 0.0001 vs. D2-EV), with p-Synapsin returning fully to control levels (Figure 3B–D).

**Figure 3.**
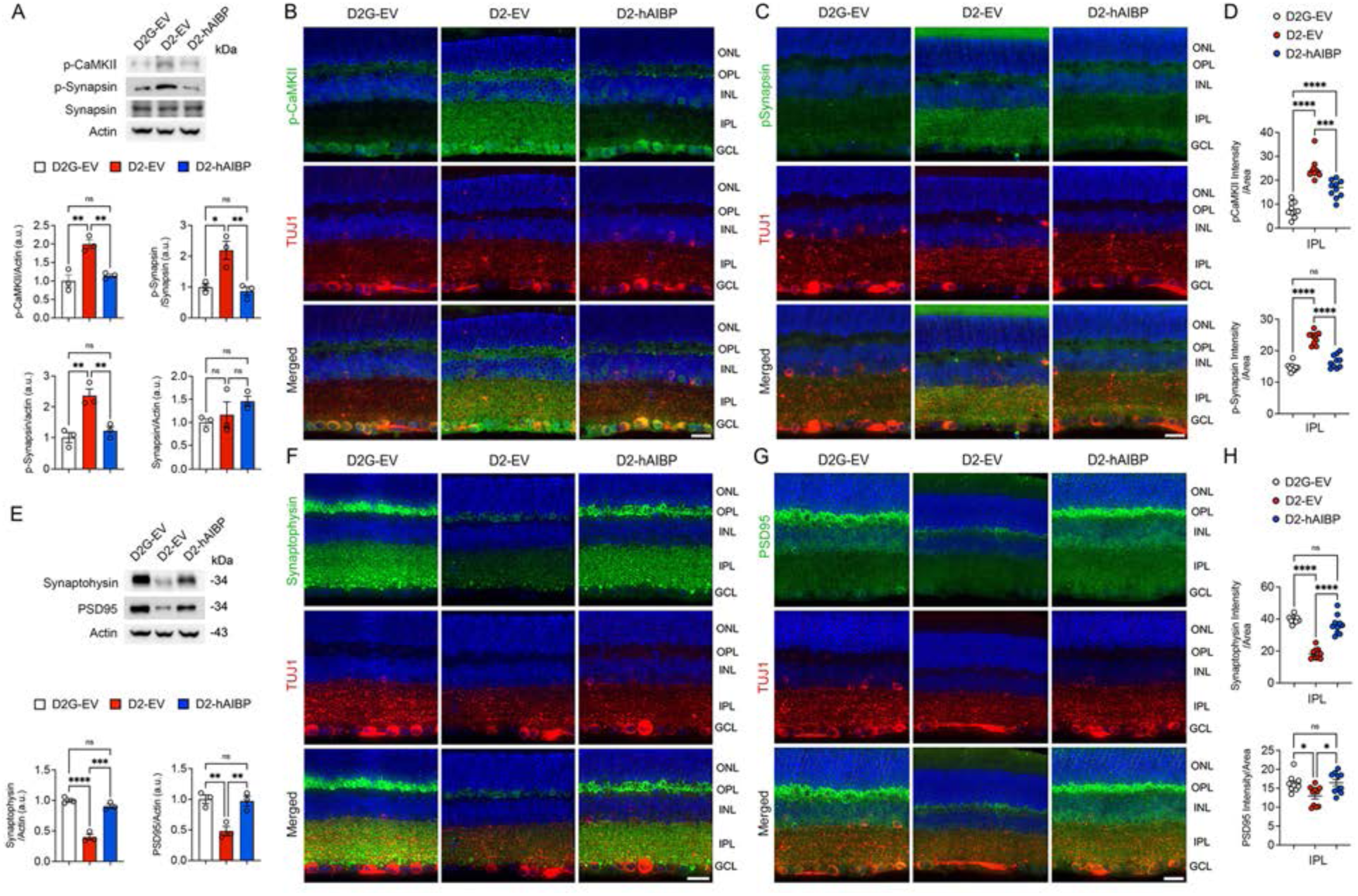
AIBP normalizes CaMKII and synapsin phosphorylation and preserves synaptic proteins in the glaucomatous D2 inner retina. (**A**) Western blot analysis of p-CaMKII, p-Synspsin, and Synapsin protein expression in the retinas of 10-month-old D2 mice (*n* = 5 mice per group). (**B**) Representative retina images for p-CaMKII (green) and TUJ1 (red) immunoreactivity. (**C**) Representative retina images for p-Synapsin (green) and TUJ1 (red) immunoreactivity. (**D**) Quantitative fluorescent intensity of p-CaMKII and p-Synapsin in the IPL of the glaucomatous retina (*n* = 5 retina sections from 3 mice per group). (**E**) Western blot analysis of synaptophysin and PSD95n protein expression in retinas from 10-month-old D2 mice (*n* = 5 mice per group). (F) Representative retina images for synaptophysin (green) and TUJ1 (red) immunoreactivity. (**G**) Representative retina images for PSD95 (green) and TUJ1 (red) immunoreactivity. (**H**) Quantitative fluorescent intensity of synaptophysin and PSD95 in the IPL of the glaucomatous retina (*n* = 5 retina sections from 3 mice per group). Error bars represent SEM. Statistical analysis was performed using one-way ANOVA and Tukey’s multiple comparisons test. \**P* < 0.05, \*\**P* < 0.01, \*\*\**P* < 0.001, and \*\*\*\**P* < 0.0001. Scale bars, 20 μm.

In parallel, the expression levels of presynaptic vesicle protein synaptophysin and the postsynaptic scaffold protein PSD95 were significantly reduced in glaucomatous D2-EV retinas by immunoblot (synaptophysin, *p* < 0.0001; PSD95, *p* < 0.01 vs. D2G-EV), but both were restored by hAIBP administration to levels not significantly different from D2G-EV controls (*p* < 0.001 and *p* < 0.01 vs. D2-EV; Figure 3E). Immunohistochemical analysis confirmed a selective loss of synaptophysin and PSD95 signal within the IPL of D2-EV retinas and its rescue in D2-hAIBP retinas (*p* < 0.0001 and *p* < 0.05 vs. D2-EV; both not significantly different from D2G-EV; Figure 3F–H). Together, these findings indicate that AIBP restrains pathological CaMKII–synapsin signaling and protects the pre- and postsynaptic machinery of the inner retina in chronic glaucoma.

### Preventive AIBP expression protects RGCs and preserves visual function in a mouse model of MB-OHT

Based on our findings of mouse AIBP-mediated neuroprotection and vision restoration in another mouse model of glaucoma,^7^ microbead (MB)-induced OHT,^7,35,36^ we injected AAV2-EV or AAV2-hAIBP intravitreally into C57BL/6 mice three weeks before MB injection (preventive mode), and followed animals for 8 weeks (Figure 4A). MB injection produced sustained IOP elevation of comparable magnitude in EV-MB and hAIBP-MB eyes throughout the 8 weeks, whereas EV-CNT and hAIBP-CNT eyes remained normotensive (Figure 4B; Table S4), suggesting that there was no IOP-lowering effect in the anterior chamber by hAIBP administration. Despite equivalent pressure exposure, RGC density in the middle and peripheral retina was significantly higher in hAIBP-MB than in EV-MB eyes (*p* < 0.001); there was no difference between normotensive EV-CNT and hAIBP-CNT eyes (Figure 4C and D; Table S4). pERG amplitude was significantly reduced by approximately 55% in EV-MB eyes (*p* < 0.0001 vs. EV-CNT) but significantly improved with hAIBP administration (*p* < 0.01 vs. EV-MB), though it did not fully normalize (*p* < 0.0001 vs. EV-CNT; Figure 4E). Importantly, hAIBP-CNT eyes did not differ from EV-CNT eyes in RGC density or pERG amplitude, indicating that AIBP overexpression is well tolerated and does not perturb normal retinal function.

**Figure 4.**
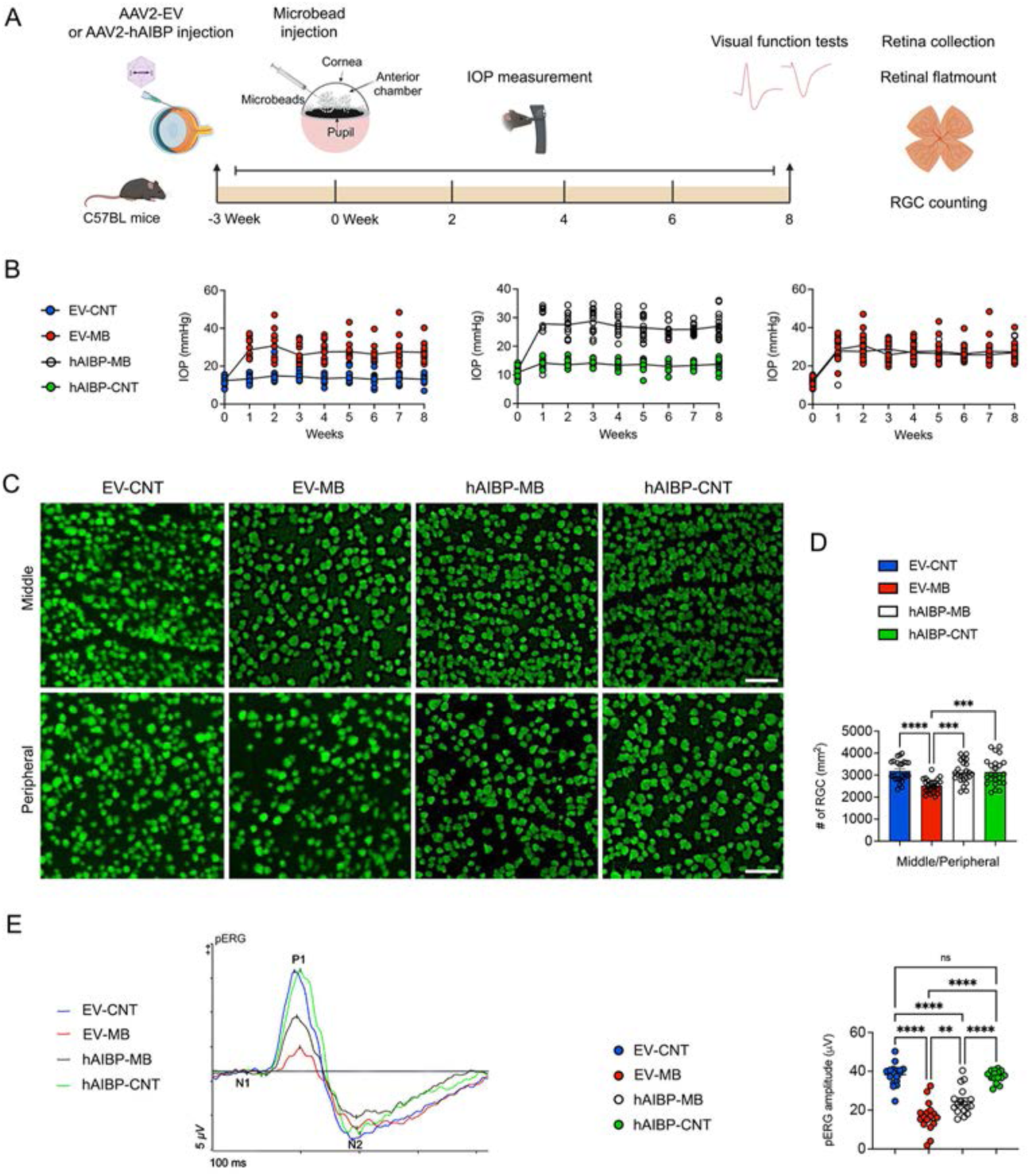
Preventive AIBP expression protects RGCs and preserves visual function in a mouse model of MB-OHT. (**A**) Experimental schematic and timeline of AAV injection, MB injection, IOP measurement, tissue collection, and data analysis in glaucomatous MB-OHT mice. (**B**) Monitoring of IOP elevation in glaucomatous MB-OHT mice (*n* = 9-11 mice per group). (**C**) Representative whole-mounted retinal images of RBPMS (green) immunoreactivity in the middle/peripheral area of the retinas. (**D**) Quantitative analysis of RGC number in the middle/peripheral areas of the retinas (*n* = 5 retina wholemounts from 5 mice for age-matched Null-Control (CNT) and AIBP-CNT mice; *n* = 7 retina wholemounts from 7 mice for Null-MB mice; *n* = 9 retinal wholemounts from 9 mice for AIBP-MB mice). (**E**) Representative recording graphs for pERG tests and quantitative analysis of pERG amplitude (*n* = 7 mice per group). Error bars represent SEM. Statistical analysis was performed using one-way ANOVA and Tukey’s multiple comparisons test. \*\**P* < 0.01, \*\*\**P* < 0.001, and \*\*\*\**P* < 0.0001. Scale bars, 50 μm. Panel (A) includes graphical elements created by BioRender (www.biorender.com).

### Therapeutic AIBP expression protects RGCs and preserves visual function in a mouse model of MB-OHT

Clinically, gene therapy would be initiated after glaucoma is diagnosed. We therefore performed the experiment with AAV2-EV or AAV2-hAIBP administered 2 weeks after MB injection, once IOP elevation was established (Figure 5A). IOP rose comparably in EV-MB and hAIBP-MB eyes and remained elevated for 8 weeks (Figure 5B; Table S5). In this therapeutic paradigm, hAIBP preserved RGC density compared to EV-MB (*p* < 0.01), to levels comparable with normotensive controls (Figure 5C and D; Table S5). pERG amplitude was significantly increased in hAIBP-MB compared to EV-MB eyes (*p* < 0.05; Figure 5E). pVEP recordings showed the clearest benefit: amplitude was restored (*p* < 0.01 vs. EV-MB) and the prolonged latency of EV-MB eyes was normalized (*p* < 0.001 vs. EV-MB), with hAIBP-MB values approaching those of EV-CNT and hAIBP-CNT eyes (Figure 5F). AIBP administration is therefore protective even when initiated after IOP elevation has begun. We next verified transgene expression and examined endogenous AIBP. His-tagged hAIBP was detected by immunoblot only in hAIBP-MB retinas (*p* < 0.001 vs. EV-CNT and EV-MB; Figure S2A) and was localized by immunostaining predominantly to TUJ1-positive RGCs in the GCL, with additional signal in the INL (*p* < 0.0001; Figure S2B and C). Notably, endogenous AIBP was significantly reduced in EV-MB retinas compared to EV-CNT (*p* < 0.05 by immunoblot; *p* < 0.0001 by IPL and GCL immunostaining), and total AIBP was restored above control levels in hAIBP-MB retinas (*p* < 0.001; Figure S2A, D, and E). OHT thus creates a state of relative AIBP deficiency in the inner retina, which AAV2-hAIBP corrects.

**Figure 5.**
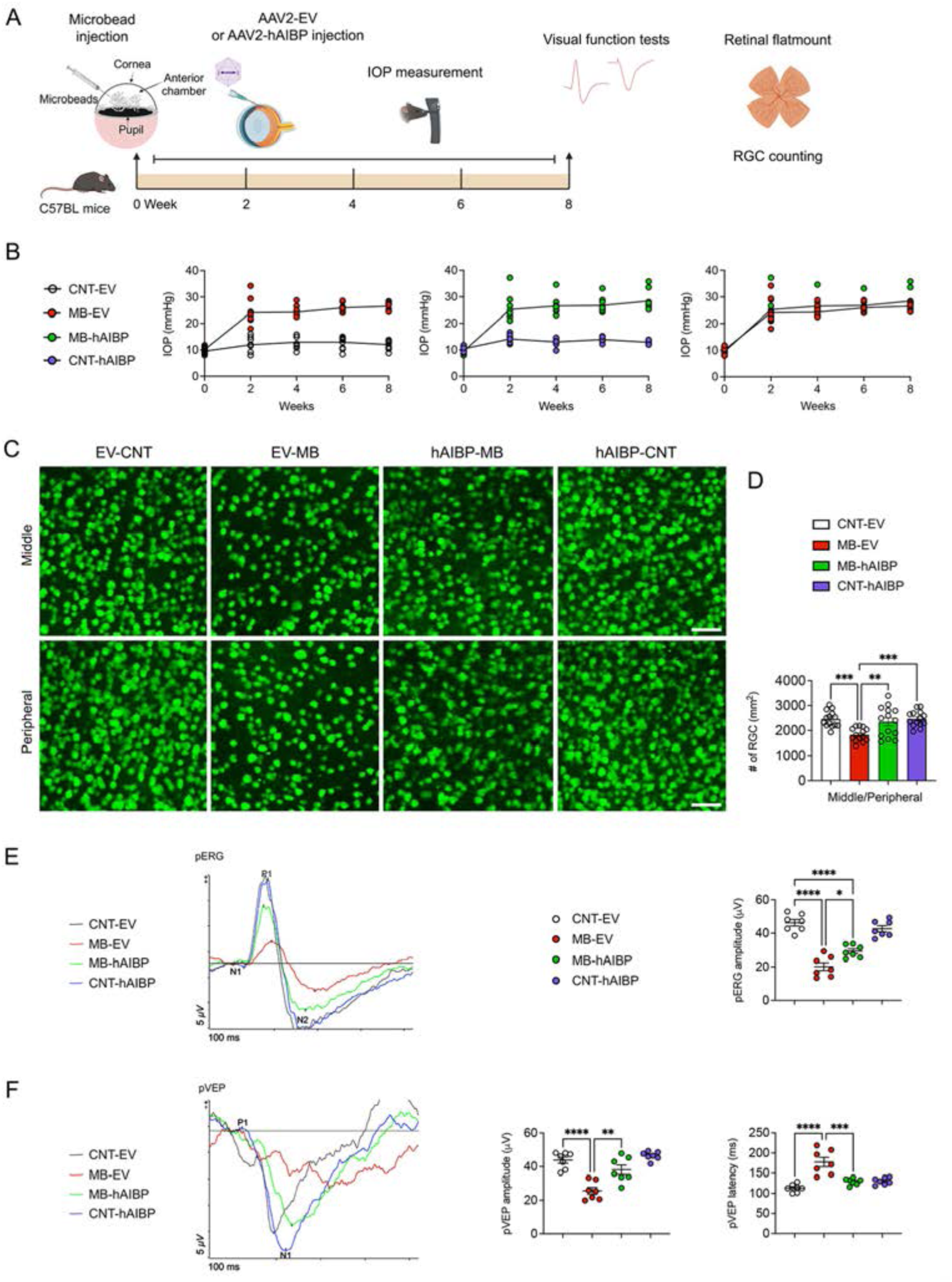
Therapeutic AIBP expression protects RGCs and preserves visual function in a mouse model of MB-OHT. (**A**) Experimental schematic and timeline of MB injection, AAV injection, IOP measurement, tissue collection, and data analysis in glaucomatous MB-OHT mice. (**B**) Monitoring of IOP elevation in glaucomatous MB-OHT mice (*n* = 9-11 mice per group). (**C**) Representative whole-mounted retinal images of RBPMS (green) immunoreactivity in the middle/peripheral area of the retinas. (**D**) Quantitative analysis of RGC number in the middle/peripheral areas of the retinas (*n* = 5 retina wholemounts from 5 mice for age-matched Null-Control (CNT) and AIBP-CNT mice; *n* = 7 retina wholemounts from 7 mice for Null-MB mice; *n* = 9 retinal wholemounts from 9 mice for AIBP-MB mice). (**E**) Representative recording graphs for pERG tests and quantitative analysis of pERG amplitude (*n* = 7 mice per group). (**F**) Representative recording graphs for pVEP tests and quantitative analysis of pVEP amplitude and latency (*n* = 7 mice per group). Error bars represent SEM. Statistical analysis was performed using one-way ANOVA and Tukey’s multiple comparisons test. \**P* < 0.015, \*\**P* < 0.01, \*\*\**P* < 0.001, and \*\*\*\**P* < 0.0001. Scale bars, 50 μm. Panel (A) includes graphical elements created by BioRender (www.biorender.com).

### AIBP restores ABCA1 expression in glaucomatous inner retina

AIBP promotes cholesterol efflux to apoA-I via ABCA1, thereby reducing plasma membrane lipid raft content and dampening TLR4 dimerization.^12,13,37^ We therefore examined ABCA1 in the inner retina. In D2 mice, ABCA1 immunoreactivity was modestly but significantly reduced in the IPL and GCL of D2-EV retinas relative to D2G-EV (*p* < 0.05 for both), and hAIBP administration increased ABCA1 above the level of both glaucomatous (*p* < 0.0001) and normotensive (*p* < 0.01) controls (Figure S3A and B). The same pattern was observed in the microbead model, where ABCA1 was markedly depleted in the IPL and GCL of EV-MB retinas (*p* < 0.0001 vs. EV-CNT) and was partially restored in hAIBP-MB retinas (IPL, *p* < 0.01; GCL, *p* < 0.05 vs. EV-MB), though not to control levels (*p* < 0.0001; Figure S3C and D). Upregulation of ABCA1 in AIBP-administered retinas is consistent with enhanced cholesterol efflux as the proximal step in AIBP-mediated suppression of TLR4 signaling.

### AIBP-mediated neuroprotection requires its TLR4-binding function in glaucomatous retina

To test whether TLR4 binding is required for protection, we generated an AAV2 vector encoding an AIBP mutant lacking the TLR4-binding domain (ΔTLR4-hAIBP). We injected AAV2-EV or AAV2-ΔTLR4-hAIBP intravitreally into C57BL/6 mice three weeks before MB injection and followed animals for 8 weeks (Figure 6A). We compared EV-CNT, EV-MB, ΔTLR4-hAIBP-MB, and ΔTLR4-hAIBP-CNT groups. IOP was elevated in EV-MB and ΔTLR4-hAIBP-MB eyes, and control eyes in both groups remained normotensive (Figure 6B; Table S6). In contrast to full-length hAIBP, ΔTLR4-hAIBP failed to preserve RGCs: RGC density in ΔTLR4-hAIBP-MB retinas was not significantly different from EV-MB in the middle retina, the peripheral retina, or the combined analysis, and remained significantly below EV-CNT in all three (*p* < 0.0001, *p* < 0.0001, and *p* < 0.001; Figure 6C and D; Table S6). Loss of the TLR4-binding domain therefore abolishes the neuroprotective activity of hAIBP, indicating that protection is mediated through the AIBP–TLR4 axis rather than through a TLR4-independent function of hAIBP. Of note, ΔTLR4-hAIBP-CNT eyes had significantly fewer RGCs than EV-CNT eyes (*p* < 0.0001; Figure 6D; Table S6), suggesting that the truncated AIBP protein may exert a dominant-negative or otherwise deleterious effect in the absence of OHT.

**Figure 6.**
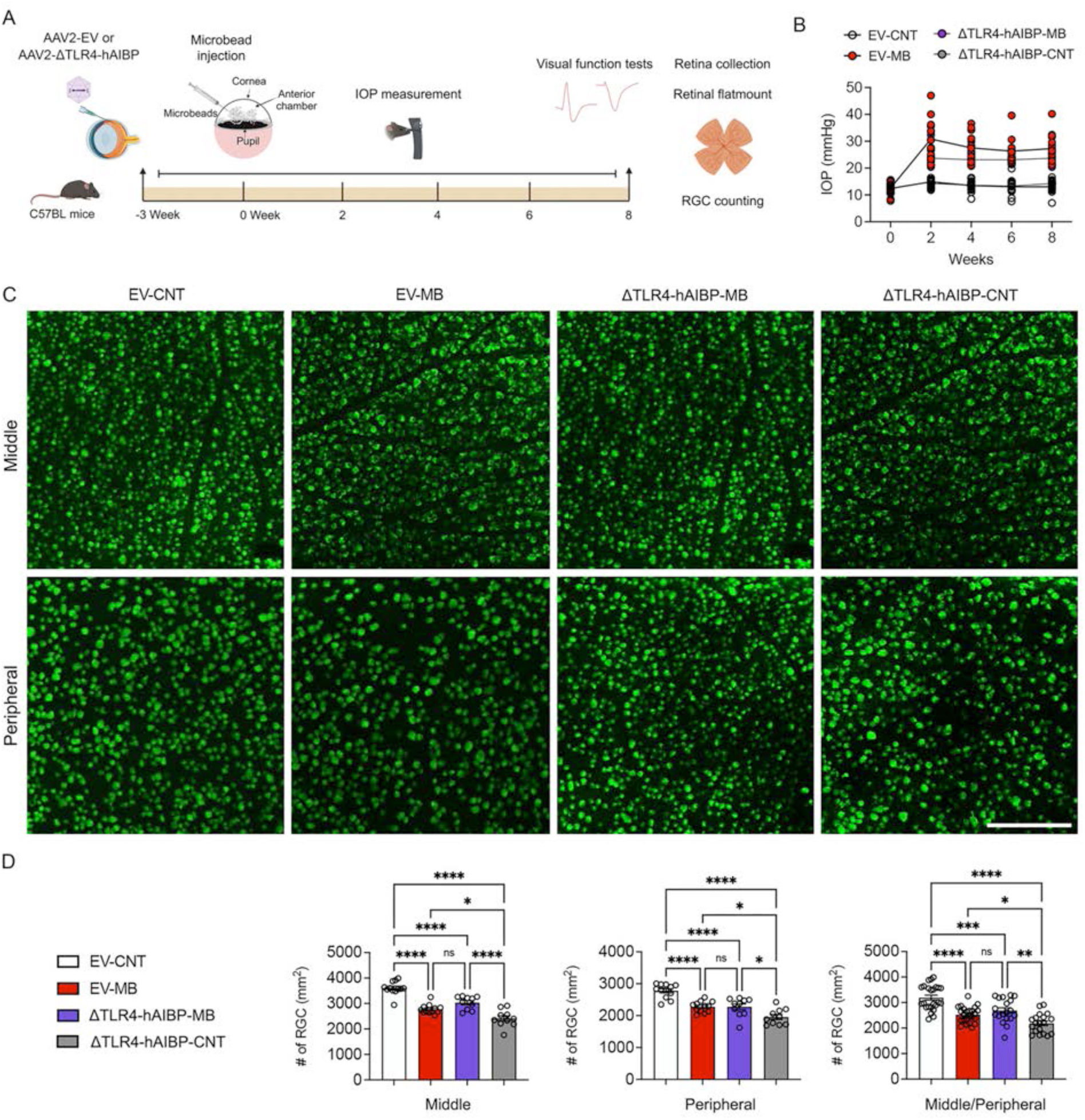
AIBP-mediated neuroprotection requires its TLR4-binding function in the MB-OHT retina. (**A**) Experimental schematic and timeline of AAV injection, MB injection, IOP measurement, tissue collection, and data analysis in glaucomatous MB-OHT mice. (B) Monitoring of IOP elevation in glaucomatous MB-OHT mice transduced with AAV2-EV or AAV2-ΔTLR4-hAIBP (*n* = 9-11 mice per group). (C) Representative whole-mounted retinal images of RBPMS (green) immunoreactivity in the middle/peripheral area of the retinas. (D) Quantitative analysis of RGC number in the middle/peripheral areas of the retinas (*n* = 10 retina wholemounts from 10 mice per group). Error bars represent SEM. Statistical analysis was performed using one-way ANOVA and Tukey’s multiple comparisons test. \**P* < 0.05, \*\**P* < 0.01, \*\*\**P* < 0.001, and \*\*\*\**P* < 0.0001. Scale bars, 50 μm. Panel (A) includes graphical elements created by BioRender (www.biorender.com).

### AIBP preserves synaptic proteins in a mouse model of MB-OHT

Consistent with the D2 data, synaptophysin and PSD95 expression were significantly reduced in EV-MB retinas by immunoblot (*p* < 0.001 and *p* < 0.01 vs. EV-CNT) but were significantly restored in hAIBP-MB retinas (both p < 0.05), to levels not significantly different from EV-CNT (Figure 7A). Immunohistochemistry showed the loss of immunoreactivity in the IPL, where synaptophysin intensity fell by approximately 60% in EV-MB eyes and was partially recovered by hAIBP (*p* < 0.05 vs. EV-MB; *p* < 0.0001 vs. EV-CNT; Figure 7B and C), and PSD95 intensity was fully restored (*p* < 0.0001 vs. EV-MB; not significant vs. EV-CNT; Figure 7D and E).

**Figure 7.**
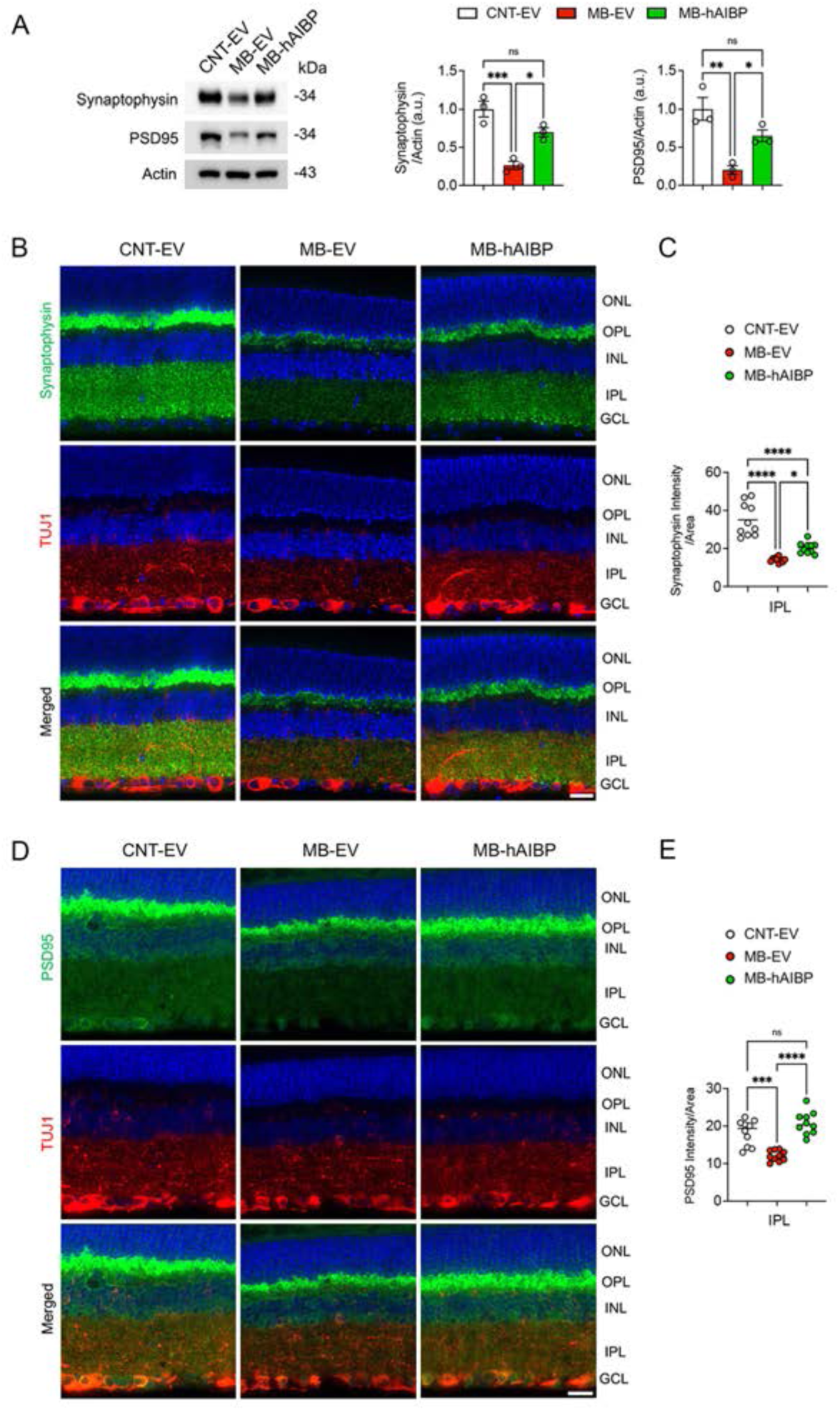
Therapeutic AIBP expression preserves synaptic proteins in a mouse model of MB-OHT. (**A**) Western blot analysis of synaptophysin and PD95 protein expression in the retinas of glaucomatous MB-OHT mice (*n* = 5 mice per group). (**B**) Representative retina images of His (green) and TUJ1 (red) immunoreactivity. (**C**) Quantitative fluorescent intensity of His in the GCL of the retinas (*n* = 5 retina sections from 3 mice per group). (**D**) Representative retina images of AIBP (green) and TUJ1 (red) immunoreactivity. (**E**) Quantitative fluorescent intensity of AIBP in the IPL and GCL of the retinas (*n* = 5 retina sections from 3 mice per group). Error bars represent SEM. Statistical analysis was performed using one-way ANOVA and Tukey’s multiple comparisons test. \**P* < 0.05, \*\**P* < 0.01, \*\*\**P* < 0.001, and \*\*\*\**P* < 0.0001. Scale bars, 20 μm.

### AIBP deficiency increases CaMKII phosphorylation and reduces synaptophysin in the retina

To determine whether endogenous AIBP is required for normal retinal synaptic homeostasis, we examined p-CaMKII and synaptophysin protein expression in the retina of wild-type (WT) and *Apoa1bp^−/−^* mice. In *Apoa1bp^−/−^* mice, immunoblotting of retinal lysates showed increased p-CaMKII in *Apoa1bp^−/−^* retinas compared to WT retinas (*p* < 0.05; Figure 8A), and immunostaining showed increased p-CaMKII in both the IPL and GCL (*p* < 0.0001; Figure 8B and C). Conversely, immunoblotting of retinal lysates showed reduced synaptophysin in *Apoa1bp^−/−^*retinas compared to WT retinas (*p* < 0.05; Figure 8D), and immunostaining showed a significant loss of IPL fluorescent signal (*p* < 0.0001; Figure 8E and F), without any experimental IOP elevation. Loss of AIBP is therefore sufficient to reproduce the CaMKII hyperphosphorylation and presynaptic protein synaptophysin loss observed in glaucomatous retinas. These findings establish AIBP as a constitutive regulator of inner retinal synaptic integrity and support a mechanistic basis for the protection conferred by AAV2-hAIBP.

**Figure 8.**
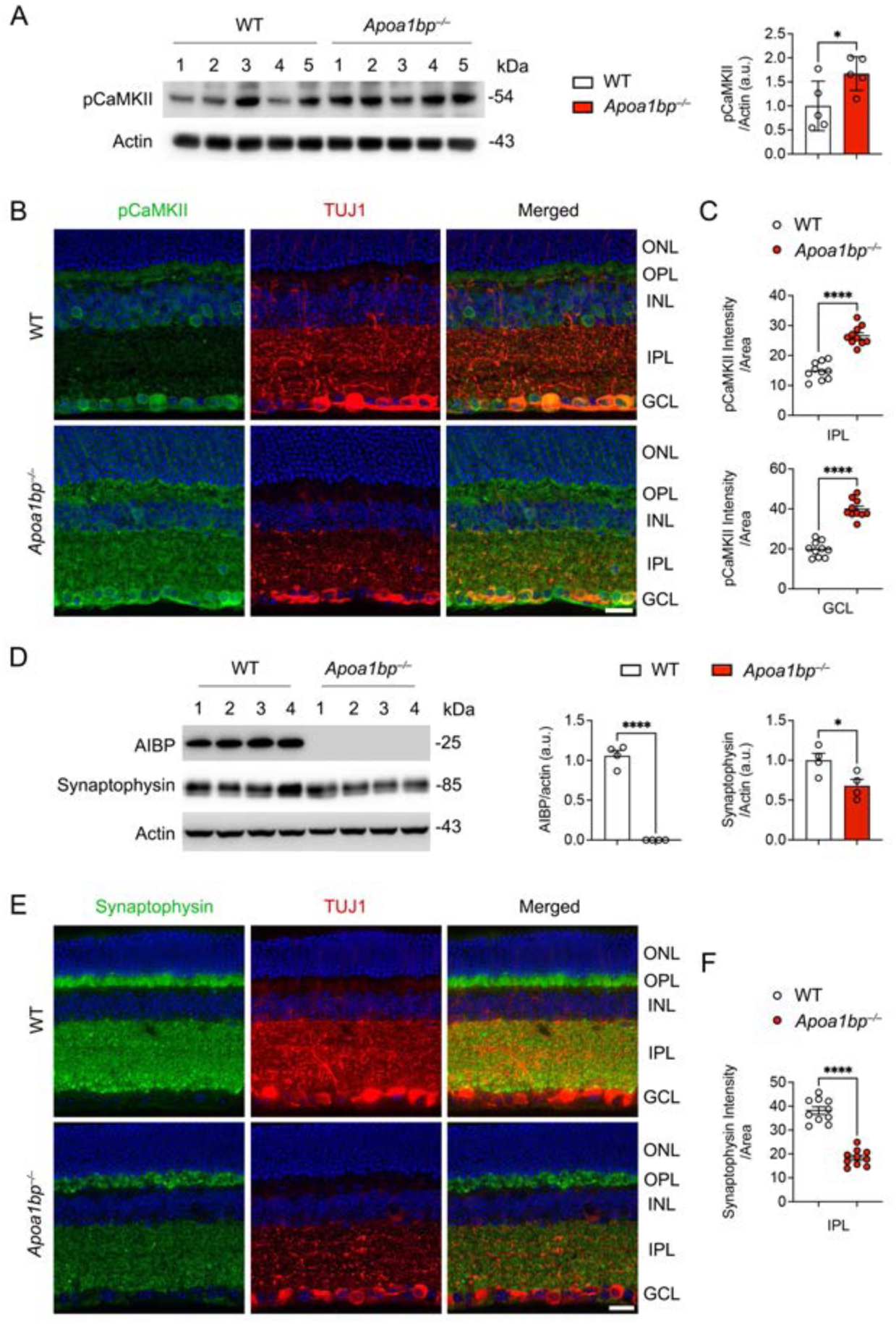
AIBP deficiency increases CaMKII phosphorylation and reduces synaptophysin in the retina. (**A**) Western blot analysis of p-CaMKII expression in the retinas of *Apoa1bp^−/−^*mice (*n* = 5 mice per group). (**B**) Representative retina images of p-CaMKII (green) and TUJ1 (red) immunoreactivity. (**C**) Quantitative fluorescent intensity of p-CaMKII in the IPL and GCL of *Apoa1bp^−/−^*retina (*n* = 5 retina sections from 3 mice per group). (**A**) Western blot analysis of AIBP and synaptophysin expression in *Apoa1bp^−/−^*retinas (*n* = 5 mice per group). (**E**) Representative retinal images of synaptophysin (green) and TUJ1 (red) immunoreactivity. (**F**) Quantitative fluorescent intensity of synaptophysin in the IPL of *Apoa1bp^−/−^*retinas (*n* = 5 retina sections from 3 mice per group). Error bars represent SEM. Statistical analysis was performed using one-way ANOVA and Tukey’s multiple comparisons test. \**P* < 0.05 and \*\*\*\**P* < 0.0001. Scale bars, 20 μm.

## DISCUSSION

The main finding of this study is that retinal AIBP overexpression, as a therapeutic approach, preserves RGC function and inner retinal synaptic activity under sustained IOP elevation. The most informative discrepancy in our data is between structural and functional endpoints. In DBA/2J mice, hAIBP rescued roughly half of the RGC loss but restored pVEP amplitude to levels statistically indistinguishable from controls. A treatment that only kept cells alive would not produce this pattern. In contrast, the pattern would be produced with a treatment that also preserved synapses and axonal signaling in surviving cells, an interpretation supported by. The synaptic data support the latter interpretation. Synaptophysin and PSD95, marking the pre- and postsynaptic compartments of the IPL, were depleted in both glaucoma models and restored by hAIBP administration, and the same pattern held in the therapeutic-delivery paradigm. This matters clinically because visual field preservation, but not RGC counts, is the goal of glaucoma treatment.

Glaucomatous retinas showed increased p-CaMKII and p-synapsin without a change in total synapsin, indicating a shift in signaling state rather than protein loss. CaMKII autophosphorylation reflects sustained Ca²⁺/calmodulin-dependent activation, and CaMKII-mediated phosphorylation of synapsin dissociates synaptic vesicles from the actin cytoskeleton, mobilizing the reserve pool^38,39^. Prolonged engagement of this axis could therefore progressively deplete presynaptic vesicle stores, consistent with the observed synaptic loss. Notably, CaMKII activity has been reported to be sensitive to metabolic stress and to promote RGC survival under certain conditions,^35,40,41^ suggesting that its effects may depend on subcellular compartment, cellular context, or duration of activation. Our data are consistent with the possibility that sustained presynaptic CaMKII signaling becomes maladaptive in the glaucomatous inner retina. Importantly, *Apoa1bp^−/−^* retinas recapitulated key features of this phenotype in the absence of OHT, including elevated p-CaMKII and reduced synaptophysin, indicating that endogenous AIBP is required to maintain inner retinal synaptic homeostasis, not merely to confer protection under glaucomatous stress. Together with our finding that OHT depletes retinal AIBP expression, this parallel finding suggests that IOP-induced AIBP deficiency may directly contribute to synaptic vulnerability rather than represent a secondary consequence of glaucomatous neuroinflammation.

The inflammatory data from this study fit the established AIBP mechanism. hAIBP increased ABCA1 in the inner retina in both models and reduced TLR4 immunoreactivity in Müller glia to control levels, consistent with enhanced cholesterol efflux depleting the lipid rafts on which TLR4 dimerization depends.^12,13^ At the downstream level, IL-1β and IL-6 expression were suppressed in Müller glia, and IBA1-positive microglial reactivity was attenuated in hAIBP-administered glaucomatous D2 retina. For a variety of reasons, C1q is of particular interest: microglia eliminate complement-tagged synapses during developmental pruning and in neurodegeneration,^22,23^ and C1q induction is among the earliest molecular events in the glaucomatous D2 retina.^4^ The concurrent reduction of C1q and preservation of synaptophysin and PSD95 in AAV2-hAIBP-administered retinas suggests that AIBP interrupts a TLR4 activation, cytokine production, complement C1q induction, and synapse elimination sequence. We emphasize that this remains correlative; direct demonstration would require, for example, synaptic engulfment assays in microglia or C1q blockade in *Apoa1bp^−/−^*retina.

The ΔTLR4-hAIBP expression provides strong evidence that TLR4 binding is necessary for retinal protection against glaucomatous neuroinflammation and neurodegeneration. Lacking the TLR4-binding domain abolished RGC protection despite equivalent IOP elevation, arguing against a purely metabolic or mitochondrial mechanism for the effect reported here. One caveat should be noted with this interpretation: ΔTLR4-hAIBP–expressing control eyes had fewer RGCs than EV controls, raising the possibility that the truncated protein is deleterious rather than simply inert, for instance by sequestering apoA-I or interfering with endogenous AIBP. Discriminating between loss-of-function and dominant-negative toxicity will require expression-matched comparisons with full-length hAIBP and direct assessment of RGC injury markers in future studies.

In this study, two important features of these data are relevant to clinical development to rescue glaucoma. First, hAIBP was protective when administered after IOP elevation was established, a realistic clinical scenario; efficacy in the post-onset paradigm with therapeutic mode was comparable to the prophylactic paradigm for RGC counts and pVEP. Second, hAIBP expression in normotensive control eyes produced no detectable change in RGC density, pERG, or pVEP compared with EV controls, consistent with AIBP’s selectivity for inflammatory and cholesterol-loaded cells.^13^ Intravitreally delivered AAV2 transduced RGCs with efficiency that increased over months, matching the chronic, decades-long course of human glaucoma, in which a single durable intervention is far preferable to repeated dosing.

The intravitreal AAV2-hAIBP restores a regulatory protein that OHT itself depletes, a deficiency we document directly in the MB-OHT model, where endogenous AIBP fell by roughly half. Our data therefore point to a mechanistically distinct route to RGC protection. Restoring an endogenously downregulated regulator may offer a conceptual advantage over supraphysiological stimulation: the pathway retains its own feedback control, and AIBP’s selectivity for cholesterol-loaded and activated cells means healthy tissue is largely spared, as indicated in the normotensive hAIBP-expressing control eyes. Whether AIBP would prove additive with metabolic or trophic strategies, or with IOP lowering, is an obvious and testable question; given the non-overlapping mechanisms, combination is a reasonable expectation rather than a remote one in future studies.

This study has several limitations. Rescue was partial for several endpoints: RGC density in glaucomatous D2 mice remained below normotensive levels, and C1q, IBA1, and IPL synaptophysin improved but did not normalize. We did not test whether higher vector doses, earlier intervention, or combination with IOP lowering would close this gap. Because the present study did not assess optic nerve axon integrity and anterograde axonal transport, we cannot dissociate the relative contributions of axonal versus synaptic preservation to the observed recovery of pVEP amplitude, and future studies will require complementary assessments (e.g., anterograde tracing, optic nerve axon counts). The glaucomatous D2 model carries pigment-dispersion-specific pathology that may not generalize, though concordant results in the MB-OHT model on a C57BL/6 background mitigate this concern. Finally, our mechanistic conclusions are based on immunohistochemical and immunoblot measures of protein expression; direct quantification of lipid raft content, TLR4 dimerization, and membrane cholesterol in this system — as has been performed previously in microglia,^13^ would provide more direct evidence for the proposed causal chain.

In conclusion, retinal hAIBP expression preserves inner retinal synaptic integrity and visual function across two distinct mouse models of glaucoma, independent of IOP reduction; this effect requires an intact TLR4-binding domain and is retained when treatment is initiated after disease onset. These findings extend the therapeutic rationale for AAV-hAIBP beyond RGC survival to preserving synaptic connectivity, which is so critical to visual function (Figure S4).

## MATERIALS AND METHODS

### Animals

Adult male and female D2 and age-matched D2-*Gpnmb^+^* mice (The Jackson Laboratory), C57BL/6J, and *Apoa1bp^−/−^* mice were housed in covered cages, fed with a standard rodent diet ad libitum, and kept on a 12 h light/12 h dark cycle. C57BL/6J mice were purchased from the Jackson Laboratory, bred in-house for experiments, and used as WT mice. *Apoa1bp^−/−^*mice on a C57BL/6J background were generated in our group, as previously reported.^42,43^ We randomly assigned animals to the experimental and control groups. To investigate the effect of AAV-hAIBP in experimental mouse models of glaucoma, we used five-month-old DBA/2J and age-matched D2-*Gpnmb^+^* mice and three-month-old WT mice.

### Production of recombinant AAV2-hAIBP vectors

AAV2-hAIBP and AAV2-ΔTLR4-hAIBP were generated as previously reported.^24^ Human AIBP (25-283aa) and ΔTLR4-hAIBP (51-283aa)^15^ were fused with FIB at the N-terminus and 6x-His at the C-terminus to generate FIB-AIBP-His and ΔTLR4-hAIBP-His, respectively, and cloned into the pAAV-MCS vector (Agilent Technologies). AAV-AIBPs were produced using the AAV2 Helper-free system (Agilent Technologies). We purified AAV2-EV, AAV2-hAIBP, and AAV2-ΔTLR4-hAIBP according to the published protocol.^44^ AAV constructs were titrated by quantitative PCR (qPCR) using primers targeting the inverted terminal repeats (ITRs) (TaKaRa Bio Inc).

### Transduction with recombinant AAV constructs

Five-month-old pre-glaucomatous D2 and D2-*Gpnmb^+^* or three-month-old C57BL/6J mice were anesthetized with an intraperitoneal (IP) injection of a mixture of ketamine/xylazine and topical 1% proparacaine eye drops. A 32-gauge needle was used to inject a total of 1 µl of AAV constructs (1 × 10^^13^ gc/ml) into the vitreous humor of the eye. Injections were administered slowly over 1 min, and the needle was kept in place for an additional 10 min to minimize vector loss through the injection tract. At the age of 10 months (D2) or 8 weeks after MB injection, the mice were euthanized by an IP injection of a mixture of ketamine/xylazine, and the retina and ONH tissues were prepared.

### Mouse model of MB-OHT

MB-OHT was induced using a modified protocol based on previous publications.^7,35,36^ In brief, eight-week-old C57BL/6J mice were intravitreally injected with either AAV-EV or AAV-AIBP as described above. At 3 weeks post-injection, mice were anesthetized by an IP injection of a cocktail of ketamine (100 mg/kg, Ketaset; Fort Dodge Animal Health) and xylazine (9 mg/kg, TranquiVed; VEDCO Inc.). We topically applied 0.5% proparacaine hydrochloride, dilated the pupils with 1% tropicamide and 2.5% phenylephrine, and then the cornea was gently punctured using a 30-gauge needle. Next, 2 µl of 1-µm-diameter polystyrene MB suspension (containing 3.0 × 10^7^ beads) followed by an air bubble, 2 µl of 6-µm-diameter polystyrene MB suspension (containing 6.3 × 10^6^ beads, Polysciences), and 1 µl of PBS containing 30% Healon was injected into the anterior chamber using a 32-gauge needle (Hamilton Company) connected to a syringe (Hamilton Company), which induced moderate elevation of IOP for ≥8 wk. An equivalent volume of PBS was injected into the contralateral eyes to serve as a sham control. At 8 weeks after the MB injection, mice were sacrificed, and the retinal tissues were prepared.

### IOP measurement

IOP elevation onset in D2 mice typically occurs between 5 and 7 months of age, and by 9 to 10 months of age, IOP-linked optic nerve axon loss is well advanced.^28,29^ We performed IOP measurements as previously described.^28,29^ Each 9- and 10-month-old D2-*Gpnmb^+^* and D2 mouse used in this study had a single IOP measurement (to confirm spontaneous IOP elevation exceeding 20 mmHg) using a tonometer (TonoLab, Colonial Medical Supply). For the MB-OHT mouse model, IOP was measured every or every other week using a tonometer (TonoLab) from 1 or 2 weeks after MB injection until 8 weeks after injection, between 3 p.m. and 5 p.m. to minimize diurnal variability. Non-IOP-elevated contralateral control retinas were used as sham control (*n* = 9-11 mice per group).

### Tissue preparation

Mice were anesthetized by an IP injection of a cocktail of ketamine/xylazine, as described above, before cervical dislocation. For immunohistochemistry, retinas were dissected from the eyeballs and fixed with 4% paraformaldehyde (Sigma) in PBS (pH 7.4) for 2 h at 4 °C. Retinas were washed several times with PBS, dehydrated through graded levels of ethanol, and embedded in polyester wax. For EM, the eyes were fixed via cardiac perfusion with 2% paraformaldehyde and 2.5% glutaraldehyde (Ted Pella) in 0.15 M sodium cacodylate (pH 7.4, Sigma) solution at 37 °C and placed in pre-cooled fixative of the same composition on ice for 1 h. For Western blot analyses, we used extracted retinas immediately.

### Western blot analyses

Harvested retinas were homogenized for 1 min on ice with a RIPA lysis buffer (Cell Signaling Technology), containing a protease inhibitor cocktail (Thermo Scientific). Proteins (10-20 μg) were run on a 4–20% Mini-PROTEAN TGX precast gels (Bio-Rad) and transferred to polyvinylidene difluoride (PVDF) membranes (GE Healthcare Bio-Science). The membranes were blocked with TBS/0.1% Tween-20 (T-TBS) containing 5% non-fat dry milk for 1 h at room temperature and incubated with primary antibodies (Supplementary Table 7) overnight at 4 °C. Membranes were washed three times with T-TBS and then incubated with horseradish peroxidase-conjugated secondary antibodies (Cell Signaling Technology) for 1 h at room temperature. Membranes were developed using an enhanced chemiluminescence substrate system (ThermoFisher Scientific). Chemiluminescent signal was detected using an enhanced chemiluminescence (ECL) substrate (Thermo Fisher Scientific) and imaged with a ChemiDoc™ Imaging System (Bio-Rad).

### Immunohistochemistry

Immunohistochemical staining of 7 μm wax sections of full-thickness retina was performed. Sections from wax blocks from each group were used for immunohistochemical analysis. To prevent non-specific background, tissues were incubated in 1% BSA/PBS for 1 h at room temperature before incubation with the primary antibodies for 16 h at 4°C. After several wash steps, the tissues were incubated with the secondary antibodies for 4 h at 4°C and washed with PBS. The sections were counterstained in PBS with the nucleic acid stain Hoechst 33342 (1 g/ml; Invitrogen). Images were acquired with Keyence All-in-One Fluorescence microscopy (BZ-X810, Keyence Corp. of America). Each target protein’s fluorescent integrated intensity in pixels per area was measured using the ImageJ software (National Institutes of Health). All imaging parameters remained the same and were corrected with background subtraction.

### Wholemount immunohistochemistry and RGC counting

Mouse retinas from enucleated eyes were dissected as flattened whole-mounts. We immersed retinas in PBS containing 30% sucrose for 24 h at 4°C. The retinas were blocked in PBS containing 3% donkey serum, 1% BSA, 1% fish gelatin, and 0.1% Triton X-100 and incubated with primary antibodies for 3 days at 4°C. We used rabbit polyclonal anti-RBPMS or mouse monoclonal anti-TUJ1 antibodies. After several wash steps, the retinas were incubated with Alexa Fluor-488-conjugated donkey anti-rabbit IgG antibody or Alexa Fluor 568-conjugated donkey anti-mouse IgG antibody for 24 h, and subsequently washed with PBS. Images were acquired with Keyence All-in-One Fluorescence microscopy (BZ-X810, Keyence Corp. of America). RBPMS- or TUJ1-positive RGCs were counted in two zones: the middle and/or peripheral retina (three-sixths and five-sixths of the retinal radius), and the scores were averaged using ImageJ.

### pERG and pVEP analyses

We measured pERG and pVEP with a Celeris apparatus (Diagnosys, MA, USA), as previously reported.^17,45^ pERG responses were recorded using alternating, reversing, black and white vertical stimuli at 1 Hz (2 reversals per second), and 50 candela/m^2^ delivered by the pattern stimulator. Then, 200 traces were recorded per eye, and averaged waveforms were calculated in which amplitudes (µV) were measured from the P1 peak to the N2 trough. At the same time, we recorded pVEP responses. For all recordings, ground and reference needle electrodes were placed subcutaneously in the tail and snout, and the active electrode was placed subdermally in the midline of the head at the location of the visual cortex. Each eye was exposed to 100 flashes of 1 Hz, 0.05 cd s/m^2^ white light through the corneal stimulators and recorded for 300 ms with a sampling frequency of 2000 Hz. Then, 200 traces were recorded per eye, and averaged waveforms were calculated in which amplitudes (µV) were measured from the P1 peak to the N1 trough. We performed five trials for each mouse and swept 100 times per trial. We set the low- and high-filter frequency cutoffs for pVEP to 1.25 Hz and 100 Hz. We analyzed the data using Espion V6 (Diagnosys).^46,47^

### Statistical analyses

All results were reported as means ± SEM. We used a two-tailed Student’s t-test to compare the two groups. For multiple group comparisons, we used one-way analysis of variance (ANOVA) and Tukey’s multiple comparisons test using GraphPad Prism. A *P* value less than 0.05 was considered statistically significant. \**P* < 0.05, \*\**P* < 0.01, \*\*\**P* < 0.001, \*\*\*\**P* < 0.0001. The value of n per group is indicated within each figure legend

## DATA AND CODE AVAILABILITY

The data presented are available in the article and the supplemental information. Additional information is available from the corresponding author upon request.

## ACKNOWLEDGEMENTS

This study was supported by National Institutes of Health grants NS129684 (WKJ), EY034116 (WKJ, SHC, KYK), AG081004 (WKJ), AG081037 (YIM, MHE, WKJ), HL171505 (YIM), P30 EY022589 (Vision core grant)

## AUTHOR CONTRIBUTIONS

Conceptualization, WKJ; Methodology, TB, SH, ZS, SHC, SC, EJ, MP, KYK, WKJ,; Investigation, TB, SH, ZS, SHC, SC, EJ, MP, KYK, WKJ; Visualization, TB, SH, ZS, KYK, WKJ; Funding acquisition, KYK, SHC, YIM, WKJ; Project administration, WKJ; Supervision, WKJ; Writing – original draft, WKJ; Writing – review & editing, TB, SHC, KYK, YIM, RNW, WKJ.

## Supplemental Information

**Figure S1.**
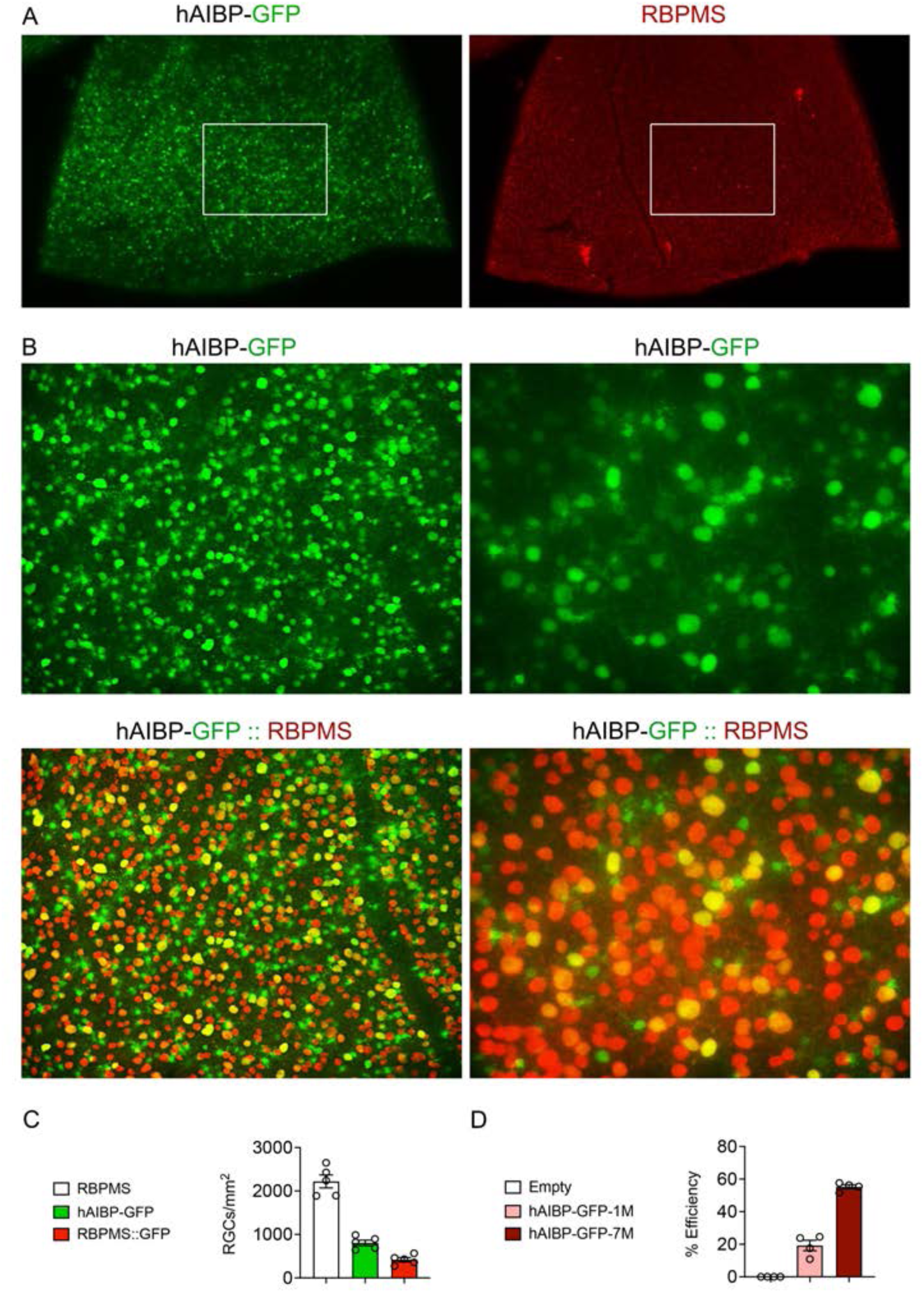
Transduction efficiency of AAV2-hAIBP-GFP in the retina. (**A**) Representative retinal wholemount images of hAIBP-GFP (green) and RBPMS (red)-positive RGCs. (**B**) Representative retinal images of hAIBP-GFP (green) and correlative GFP/RBPMS (yellow)-positive RGCs. (**C**) Quantitative analysis of RBPMS-positive, hAIBP-GFP-positive, and GFP/RBPMS-positive RGC numbers in the retina (*n* = 5 retinas from 5 mice). (**D**) Quantitative analysis of hAIBP-GFP-positive RGC numbers in the retina at 1 and 7 months following intravitreal injection of AAV2-hAIBP-GFP construct (*n* = 4 retinas from 4 mice). Scale bars, 50 μm.

**Figure S2.**
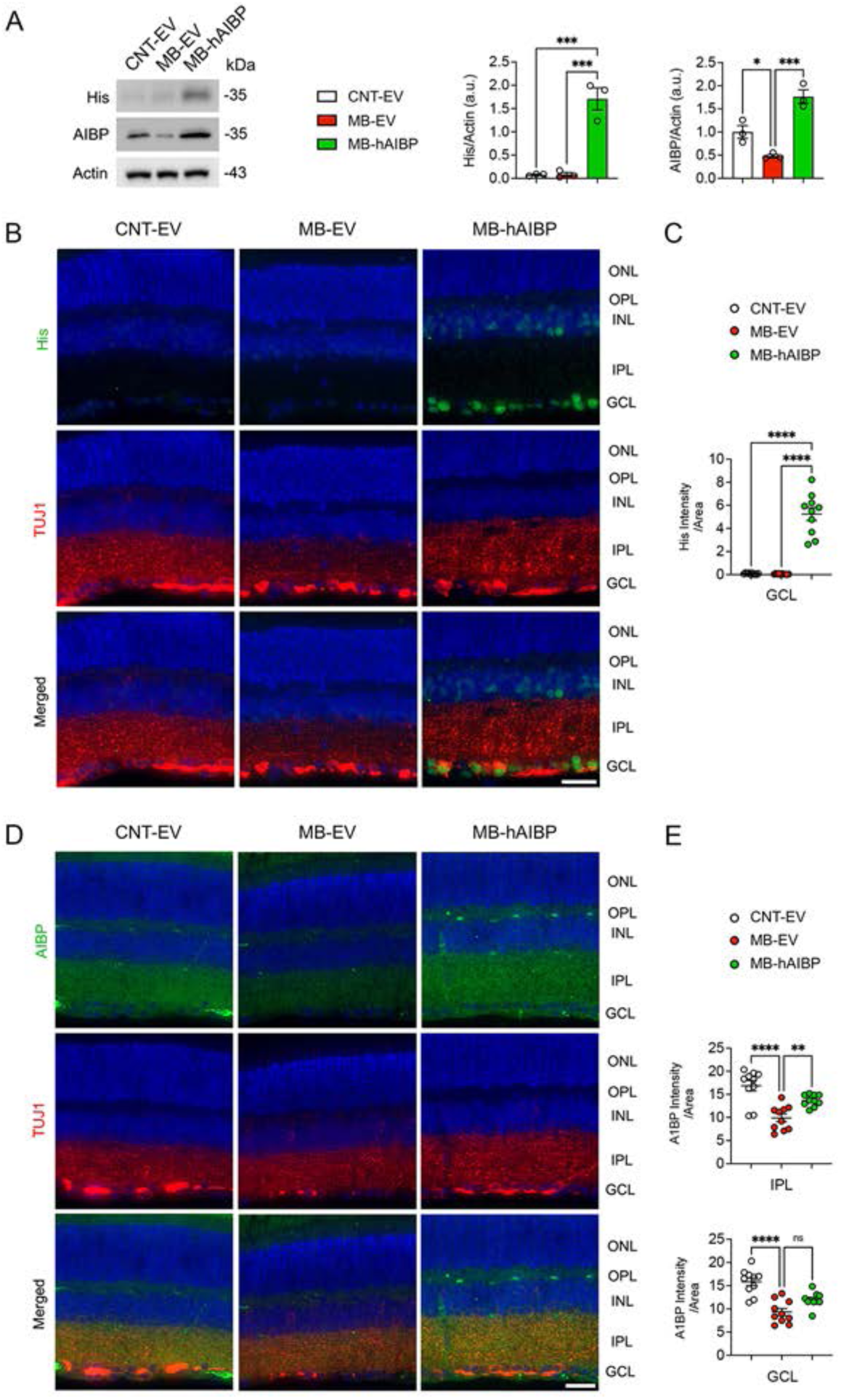
AAV2-hAIBP expression restores AIBP expression in glaucomatous retinas. (**A**) Western blot analysis of His and AIBP protein expression in the retinas of glaucomatous MB-OHT mice transduced with AAV2-EV or AAV2-hAIBP (*n* = 3 retinas from 3 mice per group). (**B**) Representative retina images of His (green) and TUJ1 (red) immunoreactivity in the retinas of glaucomatous MB-OHT mice transduced with AAV2-EV or AAV2-hAIBP. (**C**) Quantitative fluorescent intensity of His in the GCL of the glaucomatous MB-OHT mice transduced with AAV2-EV or AAV2-hAIBP (*n* = 10 retina sections from 3 mice per group). (**D**) Representative retina images of AIBP (green) and TUJ1 (red) immunoreactivity in the retinas of glaucomatous MB-OHT mice transduced with AAV2-EV or AAV2-hAIBP. (**E**) Quantitative fluorescent intensity of AIBP in the IPL and GCL of the glaucomatous MB-OHT mice transduced with AAV2-EV or AAV2-hAIBP (*n* = 10 retina sections from 3 mice per group). Error bars represent SEM. Statistical analysis was performed using one-way ANOVA and Tukey’s multiple comparisons test. \**P* < 0.05, \*\**P* < 0.01, \*\*\**P* < 0.001, and \*\*\*\**P* < 0.0001. Scale bars, 20 μm.

**Figure S3.**
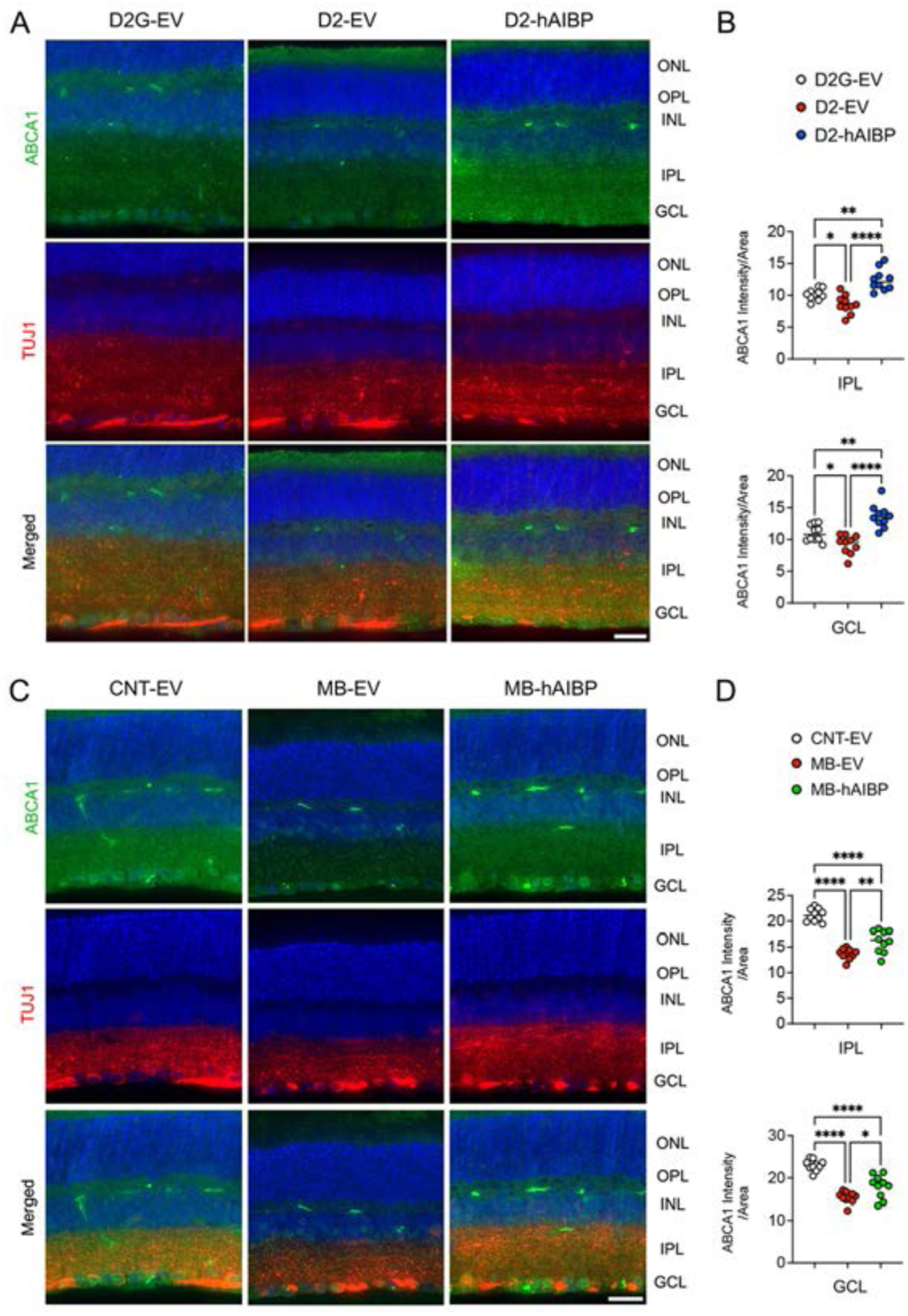
AAV2-hAIBP restores ABCA1 expression in glaucomatous retinas. (**A**) Representative retinal images of ABCA1 (green) and TUJ1 (red) immunoreactivity in glaucomatous D2 retina transduced with AAV2-EV or AAV2-hAIBP. (**B**) Quantitative fluorescent intensity of His in the IPL and GCL of the glaucomatous D2 retina transduced with AAV2-EV or AAV2-hAIBP (*n* = 10 retina sections from 3 mice per group). (**C**) Representative retina images of ABCA1 (green) and TUJ1 (red) immunoreactivity in glaucomatous MB-OHT retina transduced with AAV2-EV or AAV2-hAIBP. (**D**) Quantitative fluorescent intensity of ABCA1 in the IPL and GCL of the glaucomatous MB-OHT retina transduced with AAV2-EV or AAV2-hAIBP (*n* = 10 retina sections from 3 mice per group). Error bars represent SEM. Statistical analysis was performed using one-way ANOVA and Tukey’s multiple comparisons test. \**P* < 0.05, \*\**P* < 0.01, \*\*\**P* < 0.001, and \*\*\*\**P* < 0.0001. Scale bars, 20 μm.

**Figure S4.**
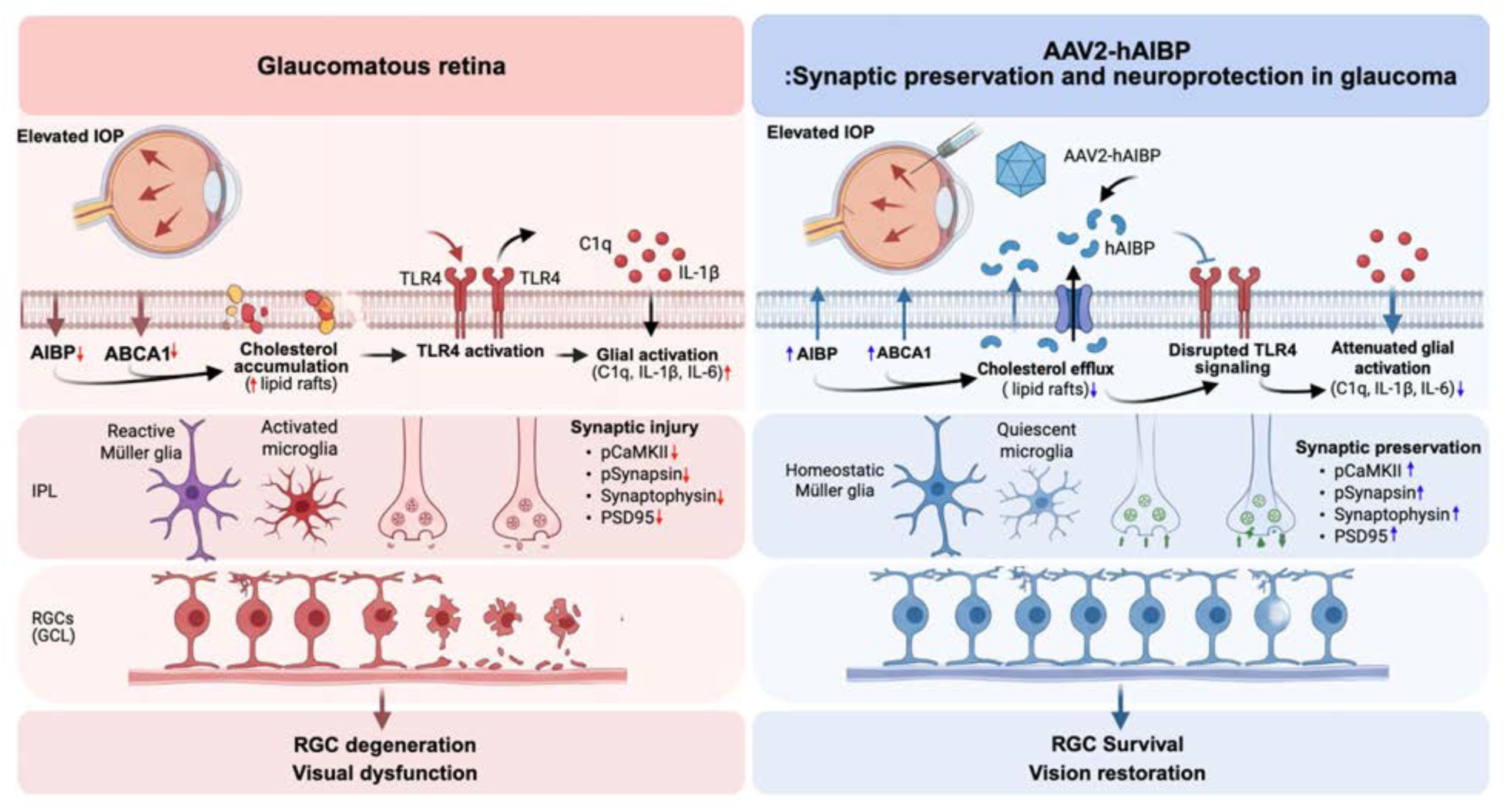
Schematic summary for the effects of hAIBP on synaptic impairment and neuroinflammation induced by glaucomatous injury. The schematic includes graphical elements created by BioRender (www.biorender.com).

## Supplementary Tables

**Supplementary Table 1.**
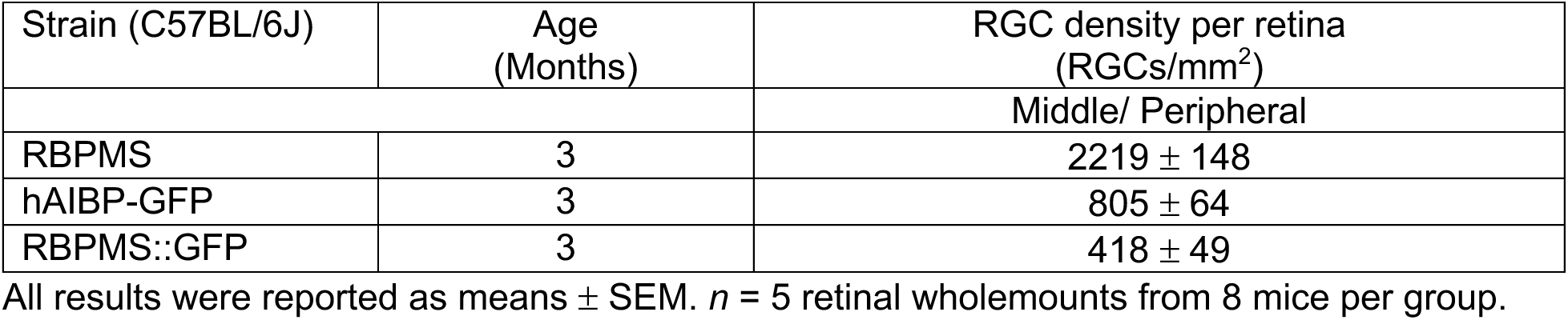
Quantitative analysis of RBPMS-positive, hAIBP-GFP-positive, and RBPMS/GFP-positive RGC numbers in the retina following intravitreal injection of AAV2-hAIBP-GFP, related to Figure S1C.

| Strain (C57BL/6J) | Age (Months) | RGC density per retina (RGCs/mm<sup>2</sup>) |
| --- | --- | --- |
|  |  | Middle/ Peripheral |
| RBPMS | 3 | 2219 ± 148 |
| hAIBP-GFP | 3 | 805 ± 64 |
| RBPMS::GFP | 3 | 418 ± 49 |

All results were reported as means ± SEM. *n* = 5 retinal wholemounts from 8 mice per group.

**Supplementary Table 2.** Quantitative analysis of hAIBP-GFP-positive RGC numbers in the retina at 1 and 7 months following intravitreal injection of AAV2-hAIBP-GFP, related to Figure S1D.

| Strain (C57BL/6J) | Age (Months) | Transduction duration (months) | Transduction efficiency (%) |
| --- | --- | --- | --- |
|  |  |  | Middle/ Peripheral |
| EV | 3 | 0 | 0 |
| hAIBP-GFP | 3 | 1 | 19.2 ± 3.1 |
| hAIBP-GFP | 3 | 7 | 55.2 ± 1.3 |

All results were reported as means ± SEM. *n* = 4 retinal wholemounts from 8 mice per group.

**Supplementary Table 3.** Effect of AAV2-hAIBP on RBPMS-positive RGC survival in the middle and peripheral retina from 10-month-old glaucomatous DBA/2J mice, related to Figure 1H-J.

| Strain (DBA/2J) | Age (Months) | Mean IOP (mmHg) at age of 10 months | Age (Months) | Mean IOP (mmHg) at age of 10 months | RGC density per retina (RGCs/mm<sup>2</sup>) |  |
| --- | --- | --- | --- | --- | --- | --- |
|  |  |  |  |  | Middle | Peripheral |
| D2G-Null | 9 | 11.4 ± 0.4 | 10 | 11.4 ± 0.4 | 3486 ± 34 | 2888 ± 65 |
| D2-Null | 9 | 24.0 ± 0.9 | 10 | 28.8 ± 1.2 | 1950 ± 168 | 1578 ± 83 |
| D2-AKAP1 | 9 | 23.4 ± 0.9 | 10 | 28.1 ± 1.2 | 2620 ± 73 | 2138 ± 74 |

All results were reported as means ± SEM. *n* = 8 retinal wholemounts from 8 mice per group.

**Supplementary Table 4.** Preventive effect of AAV2-hAIBP on RBPMS-positive RGC survival in the middle/peripheral retina from glaucomatous mice induced by MB-OHT, related to Figure 4B-D.

| Strain (C57BL/6J) | AAV Injection | Age (Months) | Mean IOP (mmHg) at 8 weeks of MB | RGC density per retina (RGCs/mm<sup>2</sup>) |
| --- | --- | --- | --- | --- |
|  |  |  |  | Middle/Peripheral |
| EV-CNT | AAV2-Null | 3 | 13.6 ± 0.22 | 3190 ± 98 |
| EV-MB | AAV2-Null | 3 | 27.7 ± 0.58 | 2524 ± 63 |
| AKAP1-MB | AAV2-AKAP1 | 3 | 27.1 ± 0.36 | 3116 ± 99 |
| AKAP1-CNT | AAV2-AKAP1 | 3 | 13.5 ± 0.15 | 3143 ± 127 |

All results were reported as means ± SEM. *n* = 7 to 9 retinal wholemounts from 7 to 9 mice per group.

**Supplementary Table 5.**
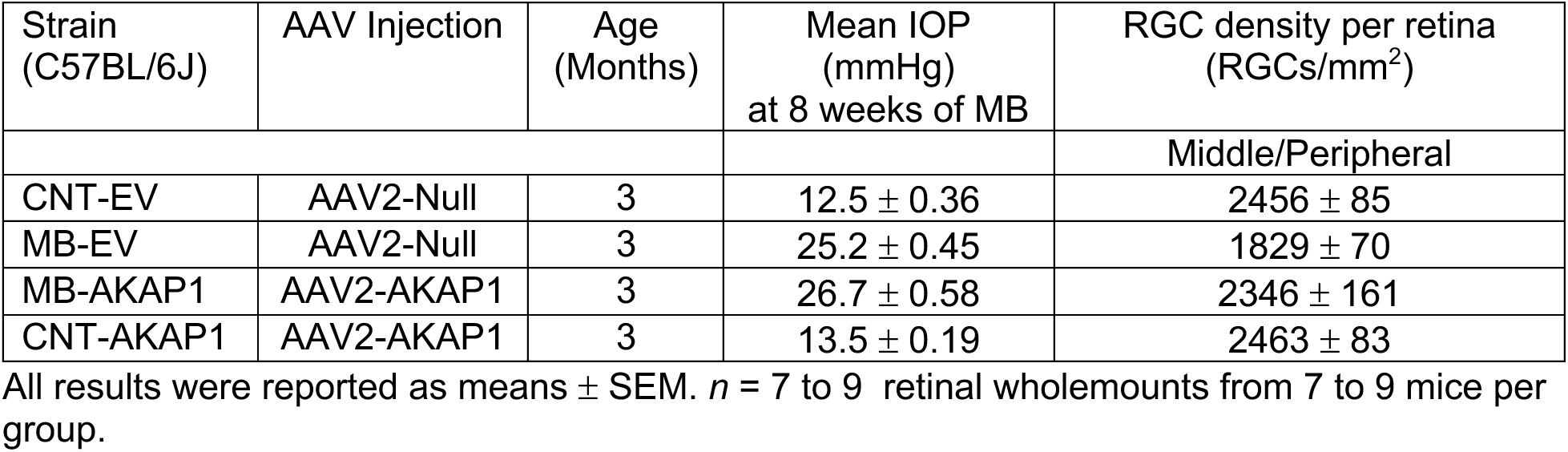
Therapeutic effect of AAV2-hAIBP on RBPMS-positive RGC survival in the middle/peripheral retina from glaucomatous mice induced by MB-OHT, related to Figure 5B-D.

**Supplementary Table 6.** Preventive effect of ΔTLR4-hAIBP on RBPMS-positive RGC survival in the middle and peripheral retina from glaucomatous mice induced by MB-OHT, related to Figure 6A-C.

| Strain (C57BL/6J) | Age (Months) | Mean IOP (mmHg) at 8 weeks of MB | RGC density per retina (RGCs/mm<sup>2</sup>) |  |
| --- | --- | --- | --- | --- |
|  |  |  | Middle | Peripheral |
| EV-CNT | 3 | 13.6 ± 0.4 | 3603 ± 77 | 2777 ± 63 |
| EV-MB | 3 | 27.3 ± 0.8 | 2774 ± 54 | 2275 ± 49 |
| ΔTLR4-hAIBP-MB |  | 23.4 ± 0.4 | 3033 ± 81 | 2273 ± 92 |
| ΔTLR4-hAIBP-CNT | 3 | 13.9 ± 0.1 | 2412 ± 105 | 1956 ± 79 |

All results were reported as means ± SEM. *n* = 10 to 12 retinal wholemounts from 10 to 12 mice per group.

**Supplementary Table 7.**
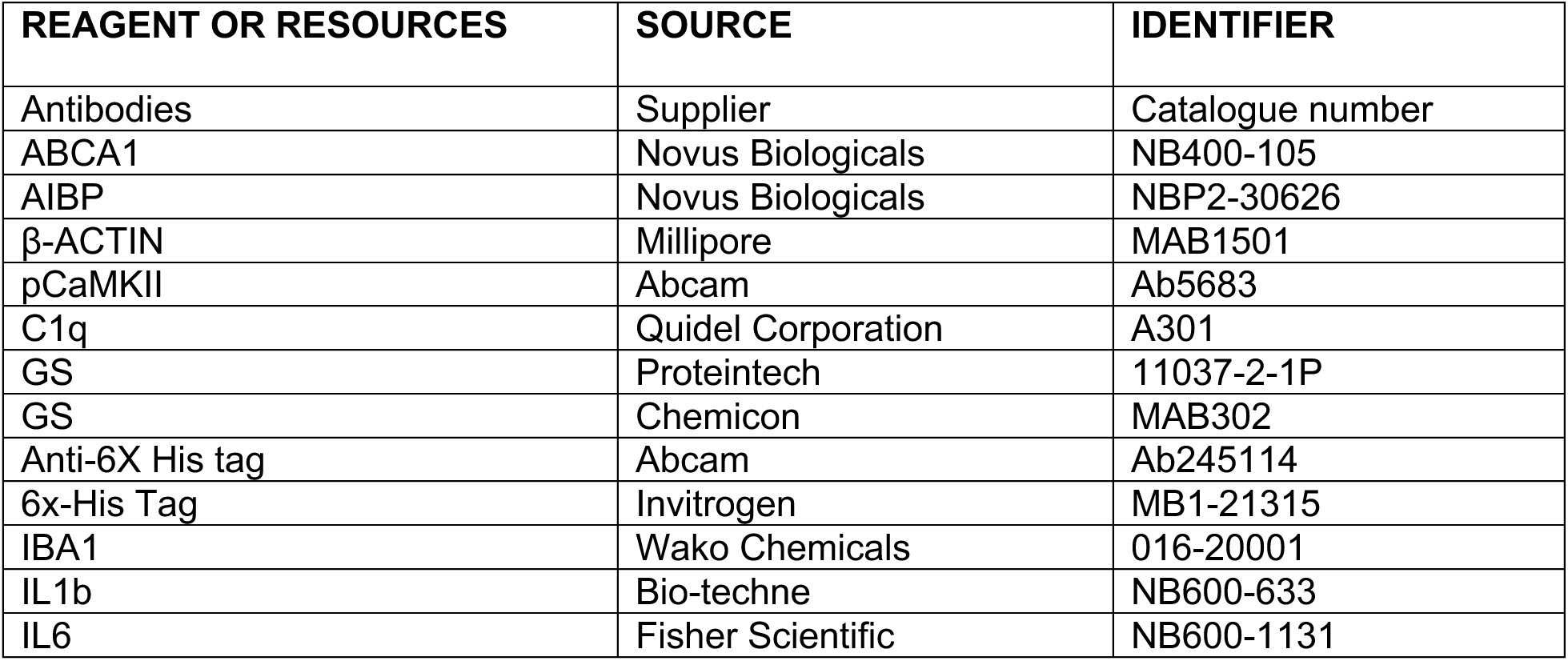
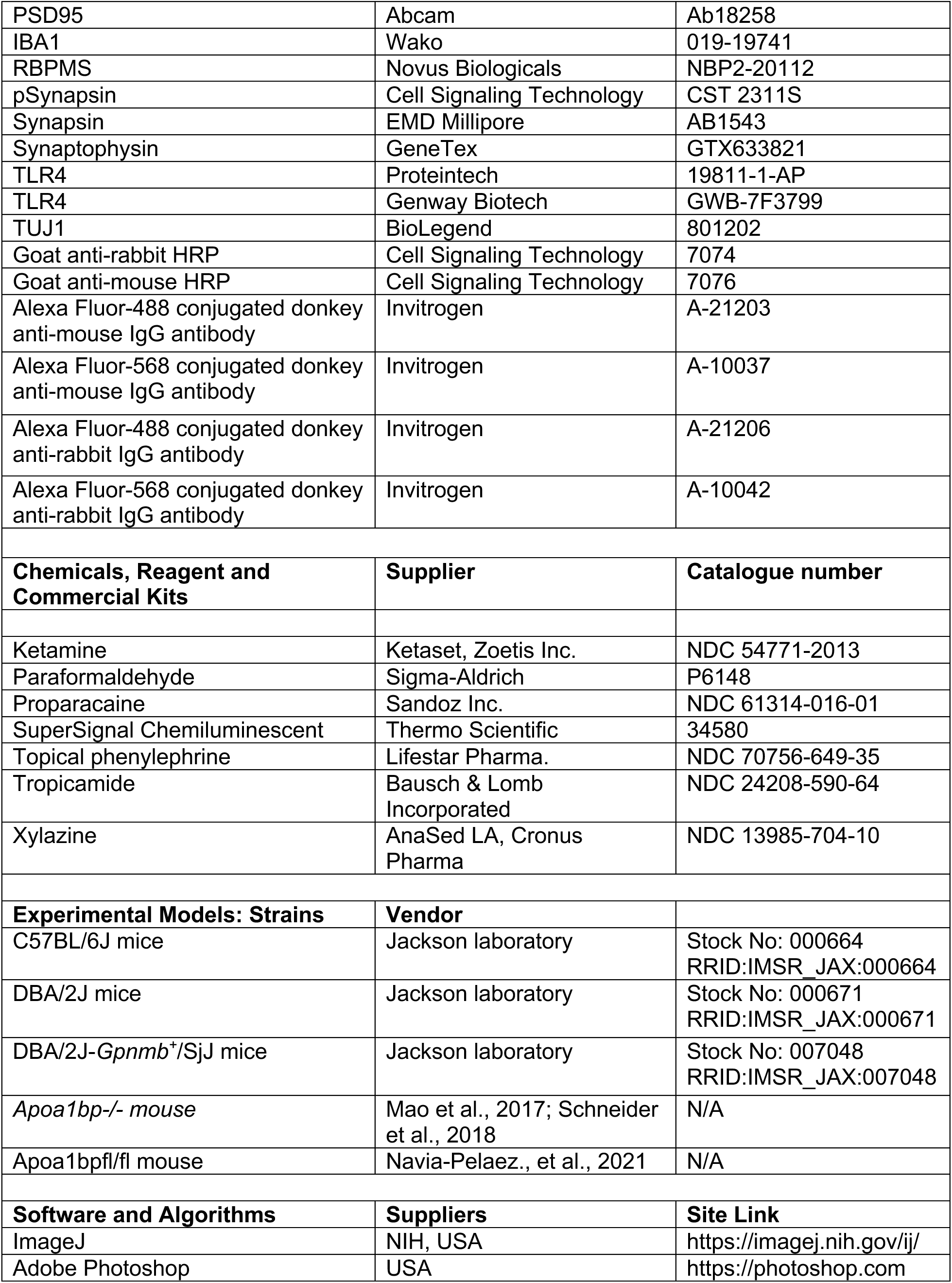
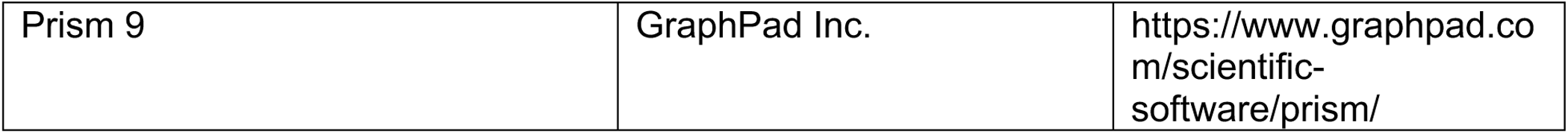
Key Resource.

